# *Drosophila insulin-like peptide 2* mediates dietary regulation of sleep intensity

**DOI:** 10.1101/681551

**Authors:** Elizabeth B. Brown, Kreesha D. Shah, Richard Faville, Benjamin Kottler, Alex C. Keene

## Abstract

Sleep is a nearly universal behavior that is regulated by diverse environmental and physiological stimuli. A defining feature of sleep is a homeostatic rebound following deprivation, where animals compensate for lost sleep by increasing sleep duration and/or sleep depth. Fruit flies exhibit robust recovery sleep following deprivation and represent a powerful model to study neural circuits regulating sleep homeostasis. Numerous neuronal populations have been identified in modulating sleep homeostasis as well as depth, raising the possibility that recovery sleep is differentially regulated by environmental or physiological processes that induce sleep deprivation. Here, we find that unlike most pharmacological and environmental manipulations commonly used to restrict sleep, starvation potently induces sleep loss without a subsequent rebound in sleep duration or depth. We find that both starvation and a sucrose-only diet result in reduced metabolic rate and increased sleep depth, suggesting that dietary yeast protein is essential for normal sleep depth and homeostasis. Finally, we find that *Drosophila insulin like peptide* 2 (*Dilp2*) is required for starvation-induced changes in sleep depth without regulating the duration of sleep. Remarkably, *Dilp2* mutant flies require rebound sleep following sleep deprivation, suggesting *Dilp2* underlies resilience to sleep loss. Together, these findings reveal innate resilience to starvation-induced sleep loss and identify distinct mechanisms that underlie starvation-induced changes in sleep duration and depth.

**Author Summary:** Sleep is nearly universal throughout the animal kingdom and homeostatic regulation represents a defining feature of sleep, where animals compensate for lost sleep by increasing sleep over subsequent time periods. Despite the robustness of this feature, surprisingly little is known about how recovery-sleep is regulated in response to different types of sleep deprivation. Fruit flies provide a powerful model for investigating the genetic regulation of sleep, and like mammals, display robust recovery sleep following deprivation. Here, we find that unlike most stimuli that suppress sleep, sleep deprivation by starvation does not require a homeostatic rebound. These findings appear to be due to flies engaging in deeper sleep during the period of partial deprivation, suggesting a natural resilience to starvation-induced sleep loss. This unique resilience to starvation-induced sleep loss is dependent on *Drosophila insulin-like peptide 2*, suggesting a critical role for insulin signaling in regulating interactions between diet and sleep homeostasis.

## Introduction

Sleep is a near universal behavior that is modulated in accordance with internal and external environments [1–3]. Numerous environmental factors can alter sleep including daily changes in light and temperature, stress, social interactions, and nutrient availability [4,5]. A central factor defining sleep is that a homeostat detects sleep loss and then compensates by increasing sleep duration and/or depth during recovery [6,7]. However, recent findings suggest that the need for recovery sleep varies depending on the neural circuits driving sleep loss [8,9]. An understanding of how different forms of sleep restriction impact sleep quality and homeostasis may offer insights into the integration of an organism’s environment with sleep drive.

Food restriction represents an ecologically-relevant perturbation that impacts sleep. In animals ranging from flies to mammals, sleep is disrupted during times of food restriction, presumably to allow for increased time to forage [10,11]. While the mechanisms underlying sleep-metabolism interactions are unclear, numerous genes have been identified that integrate sleep and metabolic state [12,13]. Further, neurons involved in sleep or wakefulness have been identified in flies and mammals that are glucose sensitive, raising the possibility that cell-autonomous nutrient sensing is critical to sleep regulation [14–16]. Despite these highly conserved interactions between sleep and metabolic regulation, little is known about the effects of starvation on sleep quality and homeostatically regulated recovery sleep when food is restored.

Fruit flies are a powerful model to study the molecular basis of sleep regulation [17,18]. Recently, significant progress has been made in identifying neural and cellular processes associated with detecting sleep debt and inducing recovery sleep [8,9,19–21]. However, a central question is how different genetic, pharmacological, and environmental manipulations differentially modulate sleep impact, quality, and homeostasis. Flies potently suppress their sleep when starved, and at least some evidence suggests they are resilient to this form of sleep loss [11,22,23]. Therefore, it is possible that starvation-induced sleep restriction is governed by neural processes that are distinct from other manipulations that restrict sleep.

Sleep quality is not only based on sleep duration, but also by sleep depth. The recent development of standardized methodology to measure sleep depth in *Drosophila* allows for investigating the physiological effects of sleep loss [24–26]. Behaviorally probing of arousal threshold and the use of indirect calorimetry to measure metabolic rate both provide indicators of sleep depth [24,25,27]. Mechanically depriving flies of sleep results in increased sleep duration and depth the following day, but the effects of other deprivation methods on sleep depth is largely unknown [24,26]. Applying these new approaches to quantify sleep during and following periods of starvation has potential to identify the mechanistic differences underlying resiliency to starvation-induced sleep loss.

Here, we find that starvation impairs sleep without inducing a homeostatic rebound. Analysis of arousal threshold reveals starved flies exhibit greater sleep depth during starvation, suggesting the lack of recovery sleep following starvation may be due to increased sleep depth during the period of food restriction. This phenotype can also be induced by feeding flies a diet lacking yeast, suggesting a critical role for dietary protein in maintaining normal sleep quality. Further, this resilience to starvation-induced sleep loss is dependent on *Drosophila insulin-like peptide 2*. Overall, these findings highlight the role of insulin signaling in regulating the interaction between diet and sleep depth.

## Results

To determine how different forms of sleep deprivation impact sleep depth and homeostatic recovery, we measured sleep and arousal threshold by video-tracking in the *Drosophila* ARousal Tracking (DART) system (Fig1A; [25]). This system probes sleep depth by measuring the responsiveness of sleeping flies to increasing intensities of mechanical stimuli (Fig 1A, representative stimulus train displayed on computer screen). After 24 hours of baseline sleep measurements in undisturbed female control (w^1118^) flies, we restricted flies of sleep for 24 hours by caffeine feeding, exposure to constant lighting, administration of mechanical stimuli, or by starvation. The sleep-restricting stimulus was then removed at ZT0 the following day to measure rebound sleep duration and arousal threshold during recovery (Fig 1B). All four manipulations significantly reduced nighttime sleep compared to undisturbed conditions (S1A and S1B Fig). As expected, sleep restricting flies through the feeding of caffeine-laced food, constant lighting, or mechanical stimulation resulted in a homeostatic rebound where flies increased sleep during the 12 hr period following sleep deprivation (Fig 1C-E, left panels). Further, these methods of sleep restriction also resulted in increased arousal threshold during rebound, indicating deeper sleep (Fig 1C-E, right panels). To ensure that this increase in arousal threshold was not a result of habituation to the mechanical stimulus over the duration of the experiment, we measured arousal threshold on standard food over a 3-day period. We found no effect of time on sleep duration or arousal threshold (S2 Fig), suggesting that this increase in arousal threshold during rebound results from sleep restriction rather than circadian regulation. Conversely, neither rebound sleep nor change in arousal threshold was detected in flies sleep-deprived by starvation (Fig 1F). Together, these findings suggest starvation restricts sleep without inducing a homeostatic increase in sleep duration or depth during rebound.

**Fig 1.**
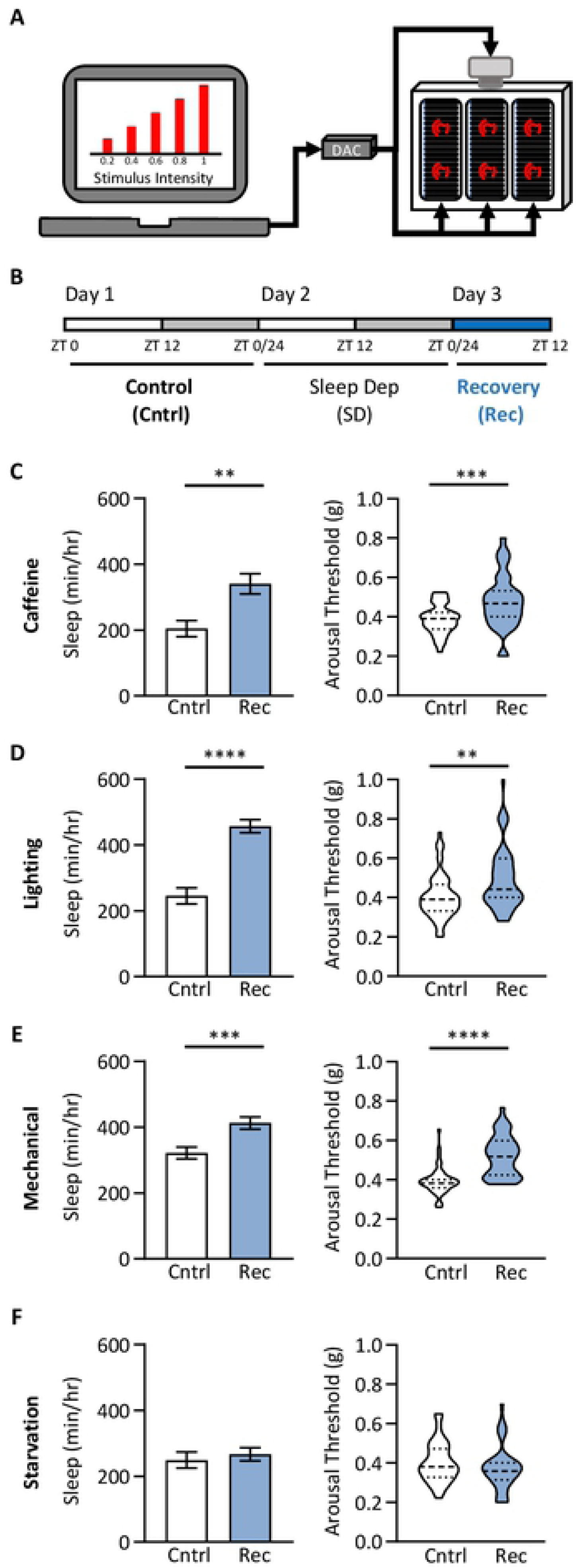
Homeostatic rebound following sleep deprivation is treatment dependent. (A) The *Drosophila* Arousal Tracking (DART) system records fly movement while simultaneously controlling mechanical stimuli via a digital analog converter (DAC). Here, mechanical stimuli are delivered to three platforms, each housing twenty flies. Mechanical stimuli of increasing strength were used to assess arousal threshold (shown on the computer screen). Arousal thresholds were determined hourly, starting at ZT0 [25]. (B) Total sleep and arousal threshold were assessed for 24 hrs on standard food (Cntrl). Flies were then sleep deprived (SD) for 24hrs using one of four treatments: 0.5mg/mL caffeine, lighting, mechanical vibration, or starvation, after which homeostatic rebound (Rec) was assessed in the subsequent 12 hrs. Comparisons were made between the first 12 hrs of the control day (white box) and the homeostatic rebound (blue box). (C-F) Daytime sleep and arousal thresholds prior to and after 24 hrs of sleep deprivation. For sleep measurements, error bars represent +/- standard error from the mean. For arousal threshold measurements, the median (dashed line) as well as 25^th^ and 75^th^ percentiles (dotted lines) are shown. (C) Sleep and arousal threshold significantly increases after 24hrs on standard food media containing 0.5mg/mL caffeine (Sleep: t-test: t_74_=3.467, *P*=0.0009; Arousal threshold: Mann-Whitney test: U=374, *P*=0.004; N = 38). (D) Sleep and arousal threshold significantly increases after 24hrs of constant lighting (Sleep: t-test: t_74_=6.782, *P*<0.0001; Arousal threshold: Mann-Whitney test: U=890.5, *P*=0.0016; N = 38). (E) Sleep and arousal threshold significantly increases after 24hrs of randomized mechanical vibration (Sleep: t-test: t_78_=3.524, *P*=0.0007; Arousal threshold: Mann-Whitney test: U=222.5, *P<*0.0001; N = 40). (F) There is no difference in sleep or arousal threshold after 24hrs of starvation (Sleep: t-test: t_70_=0.5931, *P*=0.5551; Arousal threshold: Mann-Whitney test: U=527.5, *P*=0.2402; N = 36). ** = *P*<0.01; *** = *P*<0.001; **** = *P*<0.0001.

To examine the effects of starvation-induced sleep restriction on sleep quality, we probed arousal threshold throughout the 24-hour starvation period (Fig 2A). Starvation significantly reduced sleep during both the day and night compared to flies maintained on standard fly food (Fig 2B-C). Daytime arousal threshold did not differ between fed and starved flies; yet nighttime arousal threshold was significantly increased in starved flies (Fig 2D-E). The increase in nighttime arousal threshold is specific to starvation, as there was no increase in nighttime arousal threshold when flies were sleep deprived using caffeine feeding, constant light, or mechanical vibration (S1C Fig). Together, these findings suggest flies compensate for starvation-induced sleep loss by increasing sleep depth during the night. In addition, quantifying reactivity to the mechanical stimulus as a function of individual sleep bout length revealed that arousal threshold was significantly higher in starved flies shortly after they enter sleep, suggesting starved flies are quicker to enter deeper sleep (Fig 2F).

**Fig 2.**
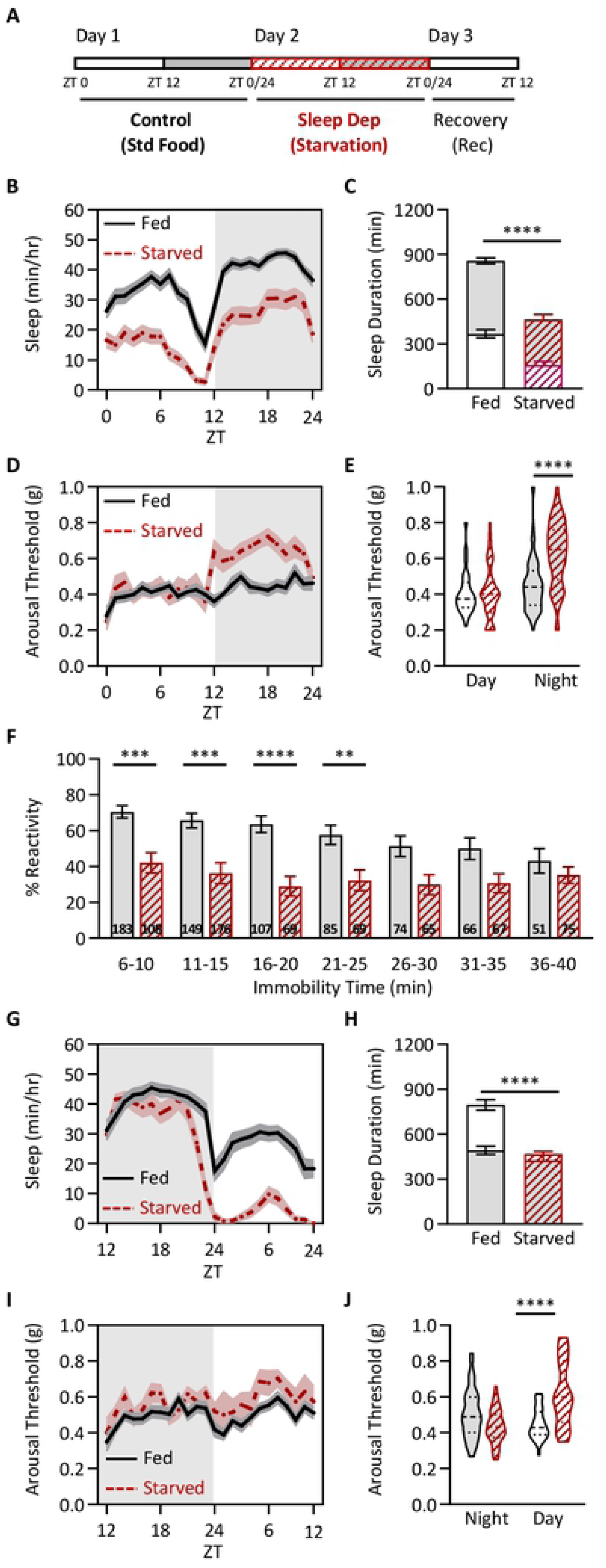
Starvation increases arousal threshold. (A) Total sleep and arousal threshold were assessed for 24 hrs on standard food (black outlined boxes) and then starved for 24hrs on agar (red outlined boxes with hatches). Flies were flipped to agar at ZT0. (B) Sleep profiles of fed and starved flies. (C) Sleep duration decreases in the starved state (two-way ANOVA: F_1,220_ = 54.63, *P*<0.0001), and occurs in both the day (*P*<0.0001) and night (*P*<0.0001). (D) Profile of arousal threshold of fed and starved flies. (E) Arousal threshold significantly increases in the starved state (REML: F_1,220_ = 34.68, *P*<0.0001), and occurs only at night (day: *P*=0.2210; night: *P*<0.0001). (F) Sleep depth is not correlated with sleep duration in starved flies. The proportion of flies that reacted to a mechanical stimulus for each bin of immobility was assessed. Starved flies are significantly less likely to respond to a mechanical stimulus during sleep (ANOVA: F_1,1230_=68.47, *P*<0.0001), specifically when flies have been sleeping for less than 30 min (6-10 min: *P*=0.0001; 11-15 min: *P*=0.0002; 16-20 min: *P*<0.0001; 21-25 min: *P*=0.0094; 26-30 min: *P*=0.0538). Numbers within each bar represent the frequency of individuals for each bin of immobility. All measurements were taken at night. (G-J) Sleep and arousal threshold measurements were taken over a 24 hr period from flies on standard food media or 1% agar. Flies were flipped to agar at ZT12. (G) Sleep profiles of fed and starved flies. (H) Sleep duration decreases in the starved state (two-way ANOVA: F_1,172_ = 32.22, *P*<0.0001), but occurs only during the day (day: *P*<0.0001; night: *P*=0.2036). (I) Profile of arousal threshold of fed and starved flies. (J) Arousal threshold significantly increases in the starved state (REML: F_1,172_ = 9.248, *P*<0.0027) and occurs only during the day (day: *P*<0.0001; night: *P*=0.0960). For sleep measurements, error bars represent +/- standard error from the mean. For arousal threshold measurements, the median (dashed line) as well as 25^th^ and 75^th^ percentiles (dotted lines) are shown. For sleep and arousal threshold profiles, shaded regions indicate +/- standard error from the mean. White background indicates daytime, while gray background indicates nighttime. ** = *P*<0.01; *** = *P*<0.001; **** = *P*<0.0001.

Sleep duration and resistance to starvation are sexually dimorphic and vary based on genetic background [28–31]. To determine whether the starvation-dependent changes in sleep depth are generalizable across sexes, we measured arousal threshold in starved male control (*w^1118^*) flies. We found that starvation results in reduced sleep duration and increased nighttime arousal threshold, phenocopying our results in females (S3A-D Fig). We next measured sleep duration and arousal threshold in Canton-S females, a second independent laboratory strain, to determine whether starvation-dependent changes in sleep depth are present in multiple laboratory strains. Again, we found that starvation decreases sleep duration and increases arousal threshold (S3E-H Fig). Overall, our findings suggest that the increase in sleep depth during starvation is generalizable across sex and *D. melanogaster* populations.

It is possible that the nighttime-specific increase in sleep depth observed in starved flies is due to either the duration of starvation or circadian regulation. To differentiate between these possibilities, we shifted the time of starvation to the onset of lights off. In these flies, there was a small reduction in nighttime sleep and a robust decrease in daytime sleep (Fig 2G,H). We found that sleep depth did not differ during the night (0-12 hrs of starvation) but was significantly increased in starved flies during the day (12-24 hrs of starvation; Fig 2I-J). Therefore, the increase in sleep depth observed in starved flies is induced by the duration of food restriction, rather than the phase of the light cycle.

Recent work suggests energy metabolism is a critical regulator of sleep homeostasis [32], but the contributions of dietary composition to sleep homeostasis remains poorly understood. Fly food is comprised primarily of sugars and protein, with yeast providing the primary protein source [33,34]. To determine the effects of different dietary macronutrients on sleep regulation, we varied the concentration of sugar and yeast and measured the effects on sleep duration and depth. Flies were fed a diet consisting of 2% yeast alone, 2% yeast and 5% sugar, or 5% sugar alone (Fig 3A). When compared to standard food, sleep duration did not differ between any of the diets, consistent with the notion that sufficiency of total caloric content, rather than the presence of a specific macronutrients regulates sleep duration (Fig 3B, S4A Fig; [11]). Quantification of arousal threshold revealed increased nighttime sleep depth in flies fed a sucrose-only diet, but not in flies fed yeast and sucrose, or yeast alone (Fig 3C, S4B Fig). These findings suggest the absence of dietary yeast increases sleep depth without affecting sleep duration.

**Fig 3.**
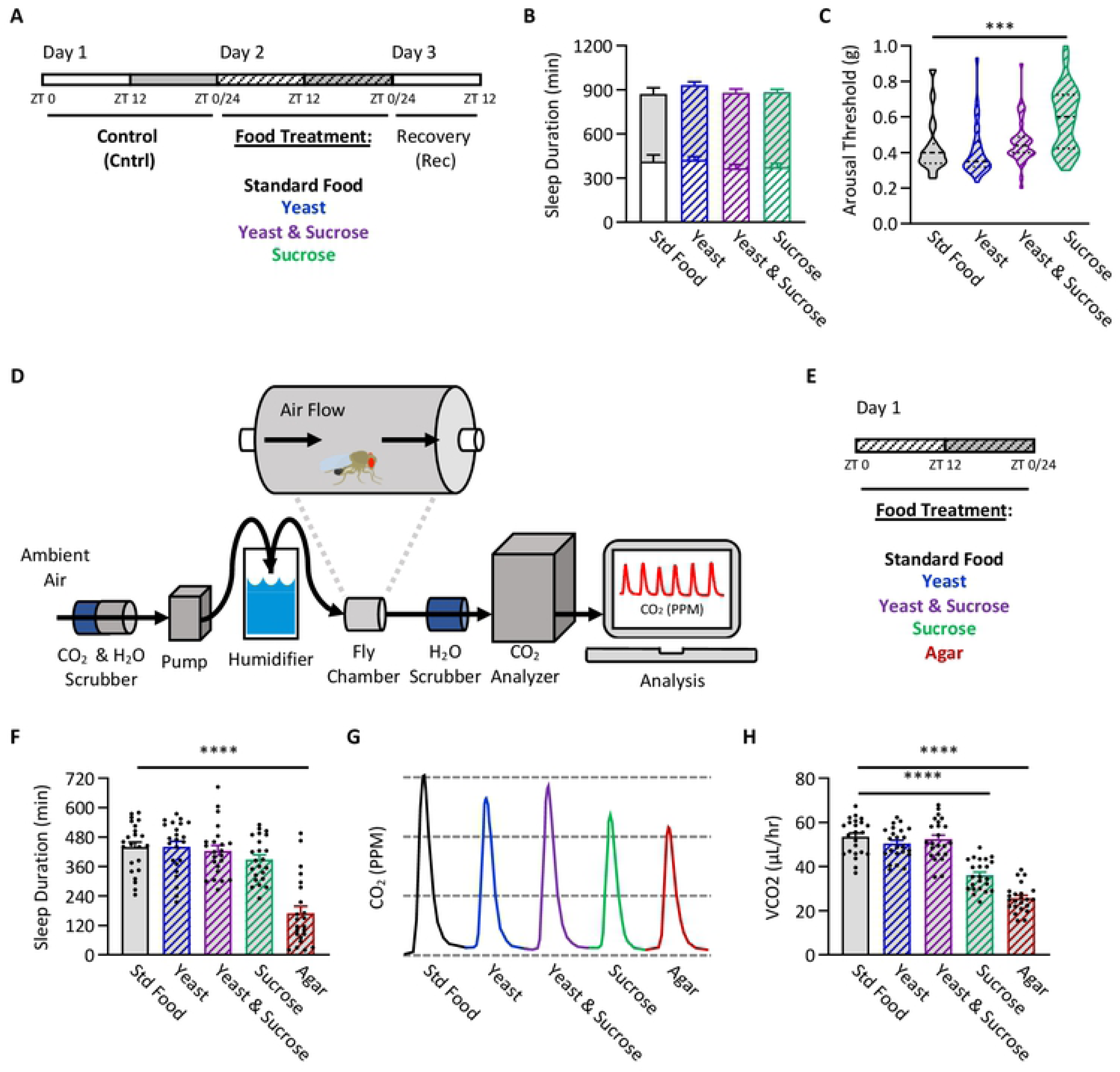
The absence of dietary yeast increases sleep depth and decreases metabolic rate. (A) Sleep and arousal threshold measurements were taken over a 24 hr period from flies fed standard food media, 2% yeast, 2% yeast and 5% sucrose, or 5% sucrose. Comparisons were then made between flies fed standard food media and each of the subsequent diets. (B) Diet does not affect sleep duration (two-way ANOVA: F_3,396_ = 0.3886, *P*=0.7612). (C) Diet does affect nighttime sleep depth (Kruskal-Wallis test: H=31.72, *P*<0.0001; N = 31-47). Post hoc analyses revealed a significant decrease in arousal threshold when flies are fed 5% sucrose, compared to standard food (*P*=0.0001). (D) Measurements of metabolic rate were taken using the SAMM system, a stop-flow respirometry system that measures the amount of CO_2_ produced over time. (E) At ZT0, adult files were placed into experimental chambers containing one of five food treatments: standard food, yeast, yeast & sucrose, sucrose, or agar. Files were allowed to acclimate to these conditions for 12hrs at which nighttime sleep duration and metabolic rate were then assessed. (F) Diet does not affect nighttime sleep duration when measured in the SAMM system, except in the absence of food (ANOVA: F_4,110_ = 25.63, *P*<0.0001, N = 23). Post hoc analyses revealed a significant decrease in sleep duration when flies are starved, in comparison to being fed standard food (*P*<0.0001). (G) Representative traces indicating the unadjusted amount of CO_2_ produced within each experimental chamber at a given timepoint for each diet. (H) Diet does affect metabolic rate (ANOVA: F_4,110_ = 59.99, *P*<0.0001, N = 24-26). Post hoc analyses revealed a significant decrease in metabolic rate when flies were fed either 5% sucrose (*P*<0.0001) or agar (*P*<0.0001), when compared to a diet of standard food. For sleep and metabolic rate measurements, error bars represent +/- standard error from the mean. For arousal threshold measurements, the median (dashed line) as well as 25^th^ and 75^th^ percentiles (dotted lines) are shown. *** = *P*<0.001; **** = *P*<0.0001.

Yeast is sensed by the taste and olfactory systems, and evidence suggests that sensory detection of yeast can influence physiology, behavior, and aging [35,36]. To determine whether differences in sensory processing contribute to the effects of diet on sleep and arousal threshold, we pharmacologically starved flies by feeding them standard fly food laced with the glycolysis inhibitor 2-deoxyglucose (2DG; S5A Fig; [37,38]). In agreement with our previous findings [38], flies fed 2DG slept less than those housed on standard food (S5B and S5C Fig). Further, this decrease in sleep duration was accompanied by an increase in arousal threshold (S5D and S5E Fig), largely phenocopying starved flies. These findings suggest that metabolic deprivation, rather than lack of sensory inputs, account for the changes in sleep duration and arousal threshold observed in starved flies.

Starved flies reduce metabolic rate, presumably to conserve energy [39], but the contributions of different dietary components on metabolic rate has not been measured in flies. To determine whether the reduction in metabolic rate in starved flies can be attributed to the absence of dietary yeast, we simultaneously measured sleep and metabolic rate in the Sleep and Activity Metabolic Monitoring (SAMM) system (Fig 3D; [39]). Consistent with data acquired in the DART system, there was no effect of diet on sleep duration in flies fed standard food, yeast, yeast and sugar, or sugar alone, and all diets resulted in more sleep than flies housed on agar (Figure 4E,F). Metabolic rate was significantly reduced in starved and sugar-fed flies compared to flies fed standard fly food or yeast diets (Figure 4G,H), suggesting that increased sleep depth, but not duration, associates with reduced metabolic rate. To determine whether the effects on metabolic rate could be recapitulated by metabolic inhibition, we next measured metabolic rate in flies fed food laced with 2DG (S6A Fig). In the SAMM system, these flies significantly reduced their nighttime sleep duration (S6B and S6C Fig). Quantification of metabolic rate revealed a significant reduction in flies fed 2DG (S6D-F Fig), thereby phenocopying results previously observed in flies fed sucrose alone. Overall, these findings suggest that both dietary protein and energy metabolism are required for normal sleep depth.

**Fig 4.**
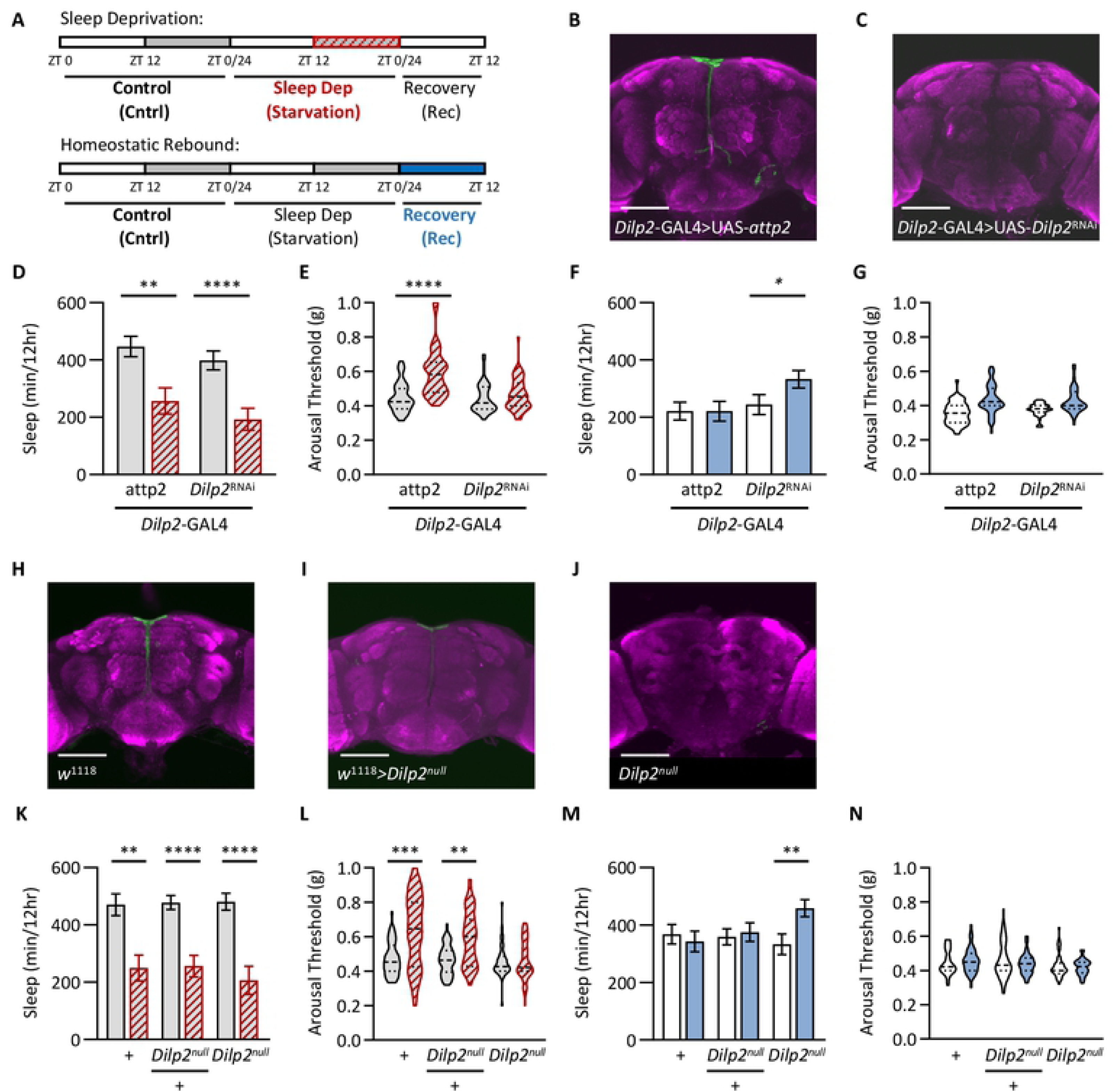
*Dilp2* uniquely regulates arousal threshold during starvation. (A) Total sleep and arousal threshold were assessed for 24 hrs on standard food (Cntrl). Flies were then sleep deprived by starving them for the following 24hrs (Sleep Dep). Homeostatic rebound (Rec) was then assessed during the subsequent 12 hrs. (B,C) Immunohistochemistry using *Dilp2* antibody (green). For each image, the brain was counterstained with the neuropil marker nc82 (magenta). Scale bar = 100μm. In comparison with the *Dilp*2-GAL4>UAS-*attp2* control (B), Dilp2 protein is reduced in *Dilp*2-GAL4>UAS-*Dilp2*^RNAi^ (C). (D) Compared to the control (*Dilp*2-GAL4>UAS-*attp2*), knockdown of *Dilp2* in *Dilp2*-expressing neurons (*Dilp*2-GAL4>UAS-*Dilp2*^RNAi^) has no effect on nighttime sleep duration (two-way ANOVA: F_1,146_ = 2.164, *P*=0.1435), however there is a significant effect of starvation (two-way ANOVA: F_1,146_ = 26.78, *P*<0.0001). For each genotype, post hoc analyses revealed a significant decrease in nighttime sleep duration when starved (*Dilp*2-GAL4>UAS-*attp2*: *P*=0.0012; *Dilp*2-GAL4>UAS-*Dilp2*^RNAi^: *P*=0.0004), when compared to standard food. (E) There is a significant effect of genotype on nighttime arousal threshold (REML: F_1,76_ = 18.22, *P*<0.0001). Post hoc analyses revealed that while controls significantly increase nighttime arousal threshold during starvation (*Dilp*2-GAL4>UAS-*attp2*: *P*<0.0001), there is no effect of knockdown of *Dilp2* in *Dilp2*-expressing neurons on arousal threshold (*Dilp*2-GAL4>UAS-*Dilp2*^RNAi^: *P*=0.5937). (F) There is a significant effect of genotype on sleep duration following 24 hrs of starvation (two-way ANOVA: F_1,146_ = 8.651, *P*=0.0038). Post hoc analyses revealed no change in daytime sleep duration in control flies (*Dilp*2-GAL4>UAS-*attp2*: *P*=0.9999), however sleep duration does significantly increase upon knockdown of *Dilp2* in *Dilp2*-expressing neurons (*Dilp*2-GAL4>UAS-*Dilp2*^RNAi^: *P*=0.0156). (G) Compared to the control (*Dilp*2-GAL4>UAS-*attp2*), knockdown of *Dilp2* in *Dilp2*-expressing neurons (*Dilp*2-GAL4>UAS-*Dilp2*^RNAi^) has no effect on arousal threshold following 24 hrs of starvation (REML: F_1,76_ = 0.4683, *P*=0.4954). (H-J) Immunohistochemistry using DILP2 antibody (green). For each image, the brain was counterstained with the neuropil marker nc82 (magenta). Scale bar = 100μm. In comparison with the w^1118^ control (H), DILP2 protein is present in heterozygotes (I), while it is absent in *Dilp2*^null^ mutants (J). (K) In comparison to the control (*w^1118^*), there is no effect on nighttime sleep duration in *Dilp2*^null^ heterozygotes or *Dilp2*^null^ mutants (two-way ANOVA: F_1,231_ = 0.1994, *P*=0.8194), however there is a significant effect of starvation (two-way ANOVA: F_1,231_ = 59.11, *P*<0.0001). For all three genotypes, post hoc analyses revealed a significant decrease in nighttime sleep duration when starved (w^1118^: *P*=0.0002; w^1118^>*Dilp2*^null^: *P*=0.0002; *Dilp2*^null^: *P*<0.0001). (L) There is a significant effect of genotype on nighttime arousal threshold (REML: F_2,117_ = 10.03, *P*<0.0001). Post hoc analyses revealed that while control flies (*w^1118^*) and *Dilp2*^null^ heterozygotes significantly increase nighttime arousal threshold during starvation (w^1118^: *P*<0.0001; w^1118^>*Dilp2*^null^: *P*<0.0001), there is no effect on arousal threshold in *Dilp2*^null^ mutants (*P*=0.9992). (M) There is a significant effect of genotype on sleep duration following 24 hrs of starvation (two-way ANOVA: F_1,231_ = 3.7810, *P*=0.0531). Post hoc analyses revealed no change in daytime sleep duration in control flies and *Dilp2* heterozygotes (*w^1118^*: *P*=0.8531; *w^1118^*>*Dilp2*^null^: *P*=0.9557), however sleep duration does significantly increase in *Dilp2*^null^ mutants (*Dilp*2^null^: *P*=0.0011). (N) In comparison to the control (*w^1118^*), there is no effect on arousal threshold following 24 hrs of starvation in *Dilp2*^null^ heterozygotes or *Dilp2*^null^ mutants (REML: F_2,117_ = 1.420, *P*=0.2347). For sleep measurements, error bars represent +/- standard error from the mean. For arousal threshold measurements, the median (dashed line) as well as 25^th^ and 75^th^ percentiles (dotted lines) are shown. ** = *P*<0.01; *** = *P*<0.001; **** = *P*<0.0001.

The *Drosophila* insulin producing cells (IPCs) are critical regulators of sleep and metabolic homeostasis [14,40,41], thereby raising the possibility that these cells regulate starvation-induced changes in sleep, arousal threshold, and metabolic rate. The IPCs express 3 of the 8 *Drosophila* insulin-like peptides (DILPS; [42,43]) and one of these, DILP2, has been previously implicated in sleep and feeding state [44,45]. To measure the effects *Dilp2* on sleep regulation, we measured sleep duration and depth in flies with disrupted *Dilp2* function (Fig 4A). We first used RNAi targeted to *Dilp2* to selectively inhibit function within the IPCs. Immunostaining with *Dilp2* antibody confirmed that *Dilp2* levels were not detectable in experimental flies (*Dilp*2-GAL4>UAS-*Dilp2*^RNAi^; Fig 4B-C). Nighttime sleep duration in flies with RNAi knockdown of *Dilp2* did not differ in the fed or starved state compared to its respective control (Fig 4D). Nighttime arousal threshold increased in starved control flies, but did not differ between fed and starved *Dilp2*-GAL4>UAS-*Dilp2*^RNAi^ flies (Fig 4E), suggesting *Dilp2* is required for starvation-dependent changes in arousal threshold. While control flies did not exhibit a rebound following deprivation, *Dilp2*-GAL4>UAS-*Dilp2*^RNAi^ flies displayed a significant sleep rebound following starvation, suggesting *Dilp2* is required for increased sleep depth during starvation that likely prevents rebound (Fig 4F). No effect on arousal threshold was observed during recovery in the control nor the *Dilp2*-GAL4>UAS-*Dilp2*^RNAi^ flies (Fig 4G). To validate our findings using RNAi, we assessed the effects of starvation on sleep depth and homeostasis in *Dilp2*^null^ flies (Fig 4H-J; [46]). Similar to RNAi knockdown, sleep duration was normal in fed and starved *Dilp2*^null^ flies, yet *Dilp2*^null^ flies did not increase arousal threshold during starvation and displayed a significant rebound in sleep upon re-feeding (Fig 4K-N). Further, during other manipulations of sleep deprivation, including feeding caffeine and constant lighting, both *Dilp2*-GAL4>UAS-*Dilp2*^RNAi^ and *Dilp2*^null^ flies respond similarly to the control in that they all decrease sleep duration with no change in depth during sleep deprivation, but following sleep deprivation produce a homeostatic rebound in which both sleep duration and depth increase (S7 Fig). Therefore, *Dilp2* is required for increasing nighttime sleep depth specifically during starvation, thereby circumventing the need for recovery sleep during the daytime when food is restored.

Given that sleep depth increases not just during starvation, but also in the absence of yeast (Fig 3C), we next tested whether *Dilp2* may similarly regulate sleep depth on a sucrose-only diet. We found that, similar to the control, feeding *Dilp2*-GAL4>UAS-*Dilp2*^RNAi^ flies a sucrose-only diet has no effect on sleep duration (S8A Fig). However, unlike the control, sleep depth does not increase on a sucrose-only diet upon knockdown of *Dilp2* using RNAi (S8B Fig). Interestingly, we did not observe any homeostatic rebound in sleep duration or depth in the control or in *Dilp2*-GAL4>UAS-*Dilp2*^RNAi^ flies (S8C and S8D Fig), suggesting that a homeostatic rebound occurs only after a decrease in both sleep duration and depth. We found similar results in *Dilp2*^null^ flies (S8E-H Fig). Overall, these results suggest that *Dilp2* uniquely regulates sleep depth both during starvation and in the absence of yeast and that the resulting homeostatic rebound is independent of sleep depth.

It is possible that *Dilp2* is required for generalized starvation-induced changes in physiology, or that it selectively regulates starvation-dependent changes in sleep depth. To differentiate between these possibilities, we measured metabolic rate in flies deficient for *Dilp2* under fed and starved conditions in the SAMM system (S9A Fig). We found no differences between control and *Dilp2*-GAL4>UAS-*Dilp2*^RNAi^ on nighttime sleep duration and metabolic rate when flies were fed standard food, sucrose only, or when starved (S9B and S9C Fig). Further, no differences in sleep and metabolic rate were detected between control and *Dilp2*^null^ flies under fed and starved conditions (S9C and S9D Fig). These findings suggest *Dilp2* is specifically required for diet dependent changes in sleep depth, but dispensable for modulation of total sleep duration and metabolic rate, and that these processes are regulated by distinct genetic architecture.

## Discussion

Animals ranging from jellyfish to humans display a homeostatic rebound following sleep deprivation [6,47]. Here we describe a functional and molecular mechanism underlying resilience to starvation-induced sleep loss. While it is widely accepted that sleep is a homeostatically regulated process, a number of factors suggest flies may be resilient to starvation-induced changes in sleep. It has previously been reported that starvation does not induce a sleep rebound [22], and in appetitive conditioning assays, starvation is required for memory formation, suggesting flies still form robust memories despite sleep loss [48–50]. Our findings suggest that changes in sleep quality during starvation underlie the reduced need for a rebound. This resiliency to starvation-induced sleep loss may reflect an ethologically relevant adaptation that allows animals to forage through portions of the night without requiring rebound sleep the following day.

A central model suggests the sleep homeostat is functionally distinct from circadian processes and detects an accumulation of sleep pressure [51]. While the cellular basis of the homeostat remains poorly understood, a large-scale screen identified the anti-microbial peptide *nemuri* as a wake-promoting factor that accumulates during periods of prolonged wakefulness [21]. In addition, numerous other genes including the TNF-alpha homolog *Eiger*, the microRNA *mir190SP*, and the rhoGTPase *Cross-veinless* are required for sleep homeostasis [52–54]. Recovery sleep following deprivation involves complex, and likely redundant neural circuits [55], raising the possibility that sleep-loss inducing perturbations can differentially impact the homeostat. For example, both the fan-shaped body and ring-neurons that comprise the ellipsoid body, brain regions associated with regulation of sleep, movement, and visual memory, have been implicated in the regulation of sleep homeostasis [9,52,56,57]. In the mushroom body, activity within a subset of sleep-promoting neurons is elevated during sleep deprivation, suggesting this may provide a neural correlate of sleep drive [58]. Further, mounting evidence suggests sleep duration can be differentiated from sleep homeostasis. As such, studies using TrpA1, the heat-activated thermosensor, to activate neurons and suppress sleep identified many neuronal populations that suppressed sleep, some of which induced a rebound while others did not [8,59]. Therefore, our findings fortify the notion that different neural processes may modulate sleep homeostasis in accordance with the perturbation that induces sleep loss.

Our findings suggest the perturbation used to deprive flies of sleep is a critical factor in regulating sleep depth and homeostasis. In fruit flies, the vast majority of studies use a ‘Sleep Nullifying Apparatus’ that consists of mechanical shaking to induce sleep deprivation in flies [60]. This method is highly effective because the timing and duration of sleep loss can be precisely controlled. Here, we find that in addition to mechanical deprivation, sleep loss induced by constant light or caffeine feeding also results in sleep rebound. Our finding that flies do not rebound after starvation-induced sleep loss can be extended to other ethologically-relevant perturbations. For example, male flies deprived of sleep by pairing with a receptive female, presumably resulting in sexual excitation, does not induce a sleep rebound when the female is removed [61,62]. Together, these findings raise the possibility that flies may be resilient to some ethologically relevant forms of sleep deprivation and highlight the importance of studying the genetic and neural processes regulating sleep across diverse environmental contexts.

In mammals, sleep stages based on cortical activity are used to measure sleep depth [63]. For example, in mammals, recovery sleep is marked by increases in slow wave sleep [64]. While orthologous sleep stages have not been identified in fruit flies, behavioral and physiological measurements suggests sleep can differ in intensity [24–26]. For example, sleep periods lasting longer than 15 minutes are associated with an elevated arousal threshold, suggesting that flies are in deeper sleep [24,25]. The finding that nighttime arousal threshold is elevated in starved animals suggests starvation enhances sleep depth. It is also is possible that arousal threshold will differ based on the sensory stimulus used to probe arousal. In agreement with this notion, it has previously been reported that starved flies are more readily aroused by odors, suggesting the olfactory system may be sensitized in starved animals [65]. Therefore, is likely that measuring arousal threshold using other sensory stimuli, including light, smell, or taste, may differentially impact arousal threshold.

While many studies have examined the interactions between sleep and feeding, these have typically compared fed and starved animals without examining the effects of specific dietary components on sleep regulation [11,36,66,67]. In *Drosophila*, feeding of a sugar only diet is sufficient for normal sleep duration [68], and evidence suggests that activation of sweet taste-receptors alone is sufficient to promote sleep [65,69]. While our group and others have interpreted this to suggest that dietary sugar alone is sufficient for ‘normal sleep’, these findings indicate that sleep quality may differ in flies fed a sugar only-diet, and that this has impacts on sleep architecture and homeostasis.

The IPCs have long been proposed as critical integrators of behavior and metabolic function [14,40,44,70]. Previous studies have implicated both the IPCs, as well as *Dilp2*, *Dilp3*, and *Dilp5* in promoting sleep during fed conditions, suggesting a role for the *Dilps* in sleep regulation [44,45]. In our analysis, we did not find any changes in sleep duration in flies lacking *Dilp2*. This is contrary to what we would predict based on previous literature, since genetic ablation of the IPCs has been shown to mimic diabetic- and starvation-like phenotypes [42,71,72]. However, possibly due to the compensatory effect of having multiple *Dilps* [46,73,74], these findings do not directly translate into expression of individual *Dilps* in the IPCs, as there are conflicting reports on whether *Dilp2* expression is modulated during starvation. Some have shown that, contrary to *Dilp3* and *Dilp5*, *Dilp2* expression does not change during starvation[42,75,76], while others have shown that *Dilp2* expression decreases [71].

Numerous factors have been identified as essential regulators of starvation-induced sleep suppression. For example, we have found that flies mutant for the mRNA/DNA binding protein *translin*, the neuropeptide *Leucokinin*, and *Astray,* a regulator of serine biosynthesis, fail to suppress sleep when starved [38,66,67]. Further, activation of orexigenic *Neuropeptide F*-expressing neurons or the sweet-sensing *Gr64f*-expressing neurons suppress sleep, suggesting that activation of feeding circuits may directly inhibit sleep [69,77]. However, it is not known whether these factors impact nutrient-dependent changes in sleep depth. A central question is how neural circuits that mediate starvation-induced sleep suppression interface with the IPCs that regulate sleep depth during the starvation period. We have found that *Leucokinin* neurons in the lateral horn signal to IPC neurons that express the *Leucokinin Receptor* to suppress sleep during starvation [67]. These findings raise the possibility of a connection between LkR signaling in the IPCs and *Dilp2* function, that in turn increase sleep depth during periods of starvation.

Overall, our findings identify differential modulation of sleep duration and depth in response to ecologically relevant sleep restriction. These findings reveal that naturally occurring resilience to starvation-induced sleep loss occurs by enhancement of sleep depth. This increase in sleep depth not only occurs during starvation, but also in the absence of yeast. Our finding that *Dilp2* is required for starvation-induced modulation of sleep depth implicates insulin signaling in its regulation and highlights the need to understand the molecular mechanisms underlying changes in sleep quality in response of environmental perturbation.

## Material and Methods

### Fly husbandry and Stocks

Flies were grown and maintained on standard *Drosophila* food media (Bloomington Recipe, Genesee Scientific, San Diego, California) in incubators (Powers Scientific, Warminster, Pennsylvania) at 25°C on a 12:12 LD cycle with humidity set to 55–65%. The following fly strains were obtained from the Bloomington Stock Center: *w^1118^* (#5905); *w^1118^*; *Dilp2^null^* (#30881; [46]); *Dilp2*-GAL4 (#37516; [72]); UAS-*attp2* (#36303; [78]); and UAS-*Dilp2* RNAi (#31068; [78]). Unless otherwise stated, 3-to-5 day old mated females were used for all experiments performed in this study.

### Sleep and Arousal Threshold Measurements

Arousal threshold was measured using the *Drosophila* Arousal Tracking system (DART), as previously described [25]. In brief, individual female flies were loaded into plastic tubes (Trikinectics, Waltham, Massachusetts) and placed onto trays containing vibrating motors. Flies were recorded continuously using a USB-webcam (QuickCam Pro 900, Logitech, Lausanne, Switzerland) with a resolution of 960×720 at 5 frames per second. The vibrational stimulus, video tracking parameters, and data analysis were performed using the DART interface developed in Matlab (MathWorks, Natick, Massachusetts). To track fly movement, raw video flies were subsampled to 1 frame per second. Fly movement, or a difference in pixilation from one frame to the next, was detected by subtracting a background image from the current frame. The background image was generated as the average of 20 randomly selected frames from a given video. Fly activity was measured as movement of greater than 3 mm. Sleep was determined by the absolute location of each fly and was measured as bouts of immobility for 5 min or more. Arousal threshold was tested with sequentially increasing vibration intensities, from 0 to 1.2 g, in 0.3 g increments, with an inter-stimulus delay of 15 s, once per hour over 24 hrs starting at ZT0.

### Sleep Deprivation

For all sleep deprivation experiments, flies were briefly anesthetized using CO_2_ and then placed into plastic tubes containing standard food. Flies were allowed to recover from CO_2_ exposure and to acclimate to the tubes for a minimum of 24 hrs prior to the start of the experiment. Upon experiment onset, baseline sleep and arousal threshold were measured starting at ZT0 for 24 hrs. For the following 24 hrs, flies were sleep deprived using one of the methods described below, during which sleep and arousal threshold were also measured. Lastly, to assess homeostatic rebound, flies were returned to standard conditions and sleep and arousal threshold was measured during the subsequent day (ZT0-ZT12). To assess the effect of sleep deprivation on sleep duration and depth, baseline nighttime sleep and arousal threshold were compared with nighttime sleep and arousal threshold measurements during sleep deprivation. To determine whether there exists a homeostatic rebound in sleep duration and/or depth, baseline daytime sleep and arousal threshold were compared to daytime sleep and arousal threshold during recovery.

#### Constant lighting

Flies were sleep-deprived for 24 hrs by maintaining constant lighting conditions during the nighttime (ZT12-24). The lighting during this time is at the same intensity as that of normal lighted conditions. During the recovery period, the lights were switched off.

#### Caffeine

Caffeine (#C0750, Sigma, St. Louis, Missouri) was added to melted standard *Drosophila* media at a concentration of 0.5 mg/mL. At ZT0 on the day of sleep deprivation, flies were transferred into tubes containing the caffeine-laced standard food media. Following 24 hrs of caffeine supplementation, flies were then transferred back into tubes containing standard *Drosophila* media during the recovery period.

#### Mechanical vibration

For mechanical vibration experiments, the motors of the DART system were utilized to disrupt sleep, as described previously [25]. Briefly, a randomized mechanical stimulus, set at a maximum of 1.2 g, occurred every 20-60 sec, with each stimulus composing 4-7 pulses lasting 0.5-4 sec in length. These randomized stimuli began at ZT0 on the day of sleep deprivation and was repeated for 24hrs.

#### Starvation

Flies were starved on 1% agar (#BP1423, Fisher Scientific, Hampton, New Hampshire), a non-nutritive substrate. At ZT0 on the day of sleep deprivation, flies were transferred into tubes containing agar. Following 24 hrs of starvation, flies were then transferred back into tubes containing standard *Drosophila* media during the recovery period.

#### Pharmacological Manipulation

2-Deoxyglucose (2DG; #D8375, Sigma) was added to melted standard *Drosophila* media at a concentration of 400 mM, as used previously to suppress sleep [38]. At ZT0 on the day of sleep deprivation, flies were transferred into tubes containing the 2DG-laced standard food media.

### Feeding Treatments

To assess sleep and arousal threshold on different diets a similar experimental design was employed as that described above. In lieu of sleep deprivation, sleep duration and depth were assessed on one of four different food treatments. Flies were anesthetized using CO_2_ and then placed into plastic tubes containing standard food. Flies were allowed to recover from CO_2_ exposure and acclimate to the tubes for a minimum of 24hrs prior to the start of the experiment. Upon experiment onset, baseline sleep and arousal threshold were measured starting at ZT0 for 24 hrs. For the following 24 hrs, flies were transferred into tubes containing one of four different diets, during which sleep and arousal threshold were again measured. These diets include standard food, 2% yeast extract (#BP1422, Fisher Scientific), 2% yeast extract and 5% sucrose (#S3, Fisher Scientific), or 5% sucrose. With the exception of standard food, all diets were dissolved in ddH2O and included 1% agar, 0.375% tegosept (methyl*-*4-hydroxybenzoate dissolved in 95% ethanol; #W271004; Sigma), and 0.125% penicillin (#15140122, ThermoFisher Scientific, Waltham, Massachusetts).

### Metabolic Rate

Metabolic rate was measured though indirect calorimetry by measuring CO_2_ production in the SAMM system, as described previously [39]. Briefly, adult flies were placed individually into behavioral chambers. Flies were acclimated to the chambers for 12hrs and then metabolic rate was assessed by quantifying the amount of CO_2_ produced in 5 min intervals during the subsequent 12hrs. For diet manipulations, each behavioral chamber contained a vial containing one of the four feeding treatments described above. For *Dilp2* experiments, each behavioral chamber contained a vial of either standard food, 5% sucrose, or 1% agar. To account for any variation between runs, each experimental run included flies from as many possible feeding treatments and/or genotypes as the system would allow.

### Immunohistochemistry

Brains of three to five day old female flies were dissected in ice-cold phosphate buffered saline (PBS) and fixed in 4% formaldehyde, PBS, and 0.5% Triton-X for 35 min at room temperature, as previously described [79]. Brains were then rinsed 3x with cold PBS and 0.5% Triton-X (PBST) for 10 min at room temperature and then overnight at 4°C. The following day, the brains were incubated for 24 hrs in primary antibody (1:20 mouse nc82; Iowa Hybridoma Bank; The Developmental Studies Hybridoma Bank, Iowa City, Iowa) diluted in 0.5% PBST at 4°C on a rotator. The brains were incubated for another 24 hrs at 4°C upon the addition of a second primary antibody (1:1300 rabbit anti-*Dilp2*). The following day, the brains were rinsed 3x in cold PBST for 10 min at room temperature and then incubated in secondary antibody (1:400 donkey anti-rabbit Alexa 488 and 1:400 donkey anti-mouse Alexa 647; ThermoFisher Scientific, Waltham, Massachusetts) for 95 min at room temperature. The brains were again rinsed 3x in cold PBST for 10 min at room temperature then stored overnight in 0.5% PBST at 4°C. Lastly, the brains were mounted in Vectashield (VECTOR Laboratories, Burlingame, California) and imaged in 2μm sections on a Nikon A1R confocal microscope (Nikon, Tokyo, Japan) using a 20X oil immersion objective. Images are presented as the Z-stack projection through the entire brain and processed using ImageJ2.

### Statistical Analysis

Measurements of sleep duration are presented as bar graphs displaying the mean ± standard error. Unless otherwise noted, a one-way or two-way analysis of variance (ANOVA) was used for comparisons between two or more genotypes and one treatment or two or more genotypes and two treatments, respectively. Measurements of arousal threshold were not normally distributed and so are presented as violin plots; indicating the median, 25^th^, and 75^th^ percentiles. The non-parametric Mann-Whitney U-test was used to compare two genotypes, while the Kruskal-Wallis test was used to compare two or more genotypes. To compare two or more genotypes and two treatments, a restricted maximum likelihood (REML) estimation was used. All post hoc analyses were performed using Sidak’s multiple comparisons test. All statistical analyses were performed using InStat software (GraphPad Software 8.0).

## Acknowledgements

We are thankful to members of the Keene laboratory for helpful discussions and technical support. We are also thankful to Jan Veenstra for kindly providing anti-*Dilp2* antibody and Dragana Rogulja for helpful discussion. This work was supported by NIH Grants R01 HL143790 and R01 NS085152 to ACK, as well as support from FAU’s Jupiter Life Science Initiative.

## Author Contributions

Conceptualization: EBB, ACK

Data Curation: EBB, KDS, RF, BK, ACK

Formal Analysis: EBB, KDS

Funding Acquisition: ACK

Investigation: EBB, KDS, ACK

Methodology: EBB, KDS, RF, BK, ACK

Project administration: EBB, ACK

Resources: EBB, RF, BK, ACK

Software: RF, BK

Supervision: EBB, ACK

Validation: EBB

Visualization: EBB

Writing – original draft preparation: EBB, ACK

Writing – review & editing: EBB, KDS, RF, BK, ACK

## Supporting Information

**S1 Fig.**
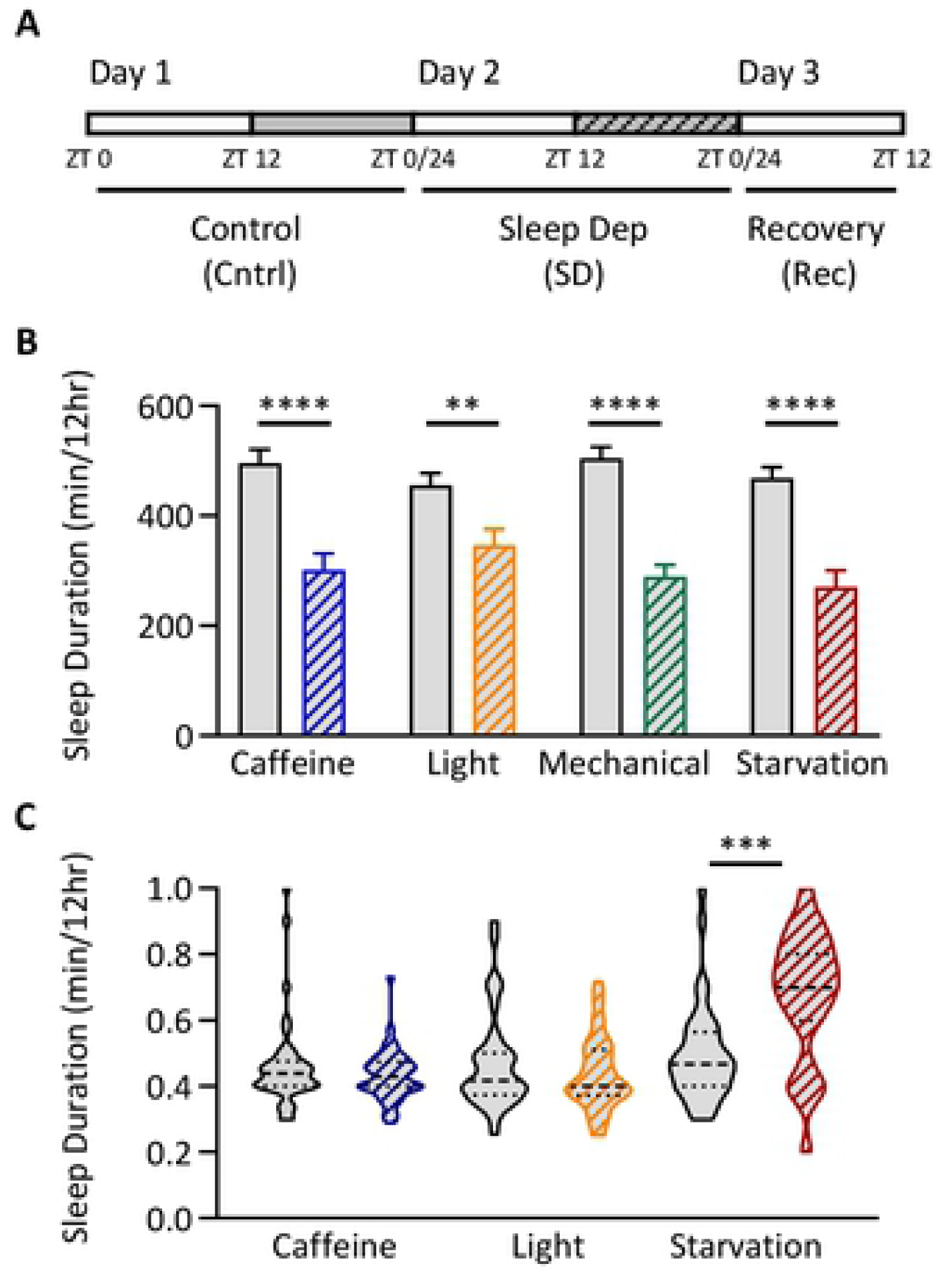
Unlike starvation, other methods of sleep deprivation do not increase nighttime arousal threshold. (A) Total sleep was assessed for 24 hrs on standard food (Cntrl). Flies were then sleep deprived for 24hrs using one of four treatments: 0.5mg/mL caffeine, constant lighting, mechanical vibration, or starvation. Comparisons were made between the control night (solid gray box) and the nighttime during sleep deprivation (black hatched box). (B) Nighttime sleep duration significantly decreases for each method of sleep deprivation (0.5mg/mL caffeine: t-test: t_78_=4.986, *P*<0.0001, constant lighting: t-test: t_78_=2.996, *P*=0.0037; mechanical vibration: t-test: t_78_=7.298, *P*<0.0001; starvation t-test: t_74_=5.495, *P*<0.0001). (C) With the exception of starvation, arousal threshold remains unchanged during the night when using each of the described methods of sleep deprivation (0.5mg/mL caffeine: U=659.5, *P*=0.5165, N = 38; constant lighting: U=617.5, *P*=0.4715, N = 37; starvation: U=225, *P*=0.0001, N = 37). Arousal threshold measurements via mechanical vibration were unable to be calculated since the mechanical vibration used to assess arousal threshold was being used to implement sleep deprivation. For sleep measurements, error bars represent +/- standard error from the mean. For arousal threshold measurements, the median (dashed line) as well as 25^th^ and 75^th^ percentiles (dotted lines) are shown. ** = *P*<0.01; *** = *P*<0.001; **** = *P*<0.0001.

**S2 Fig.**
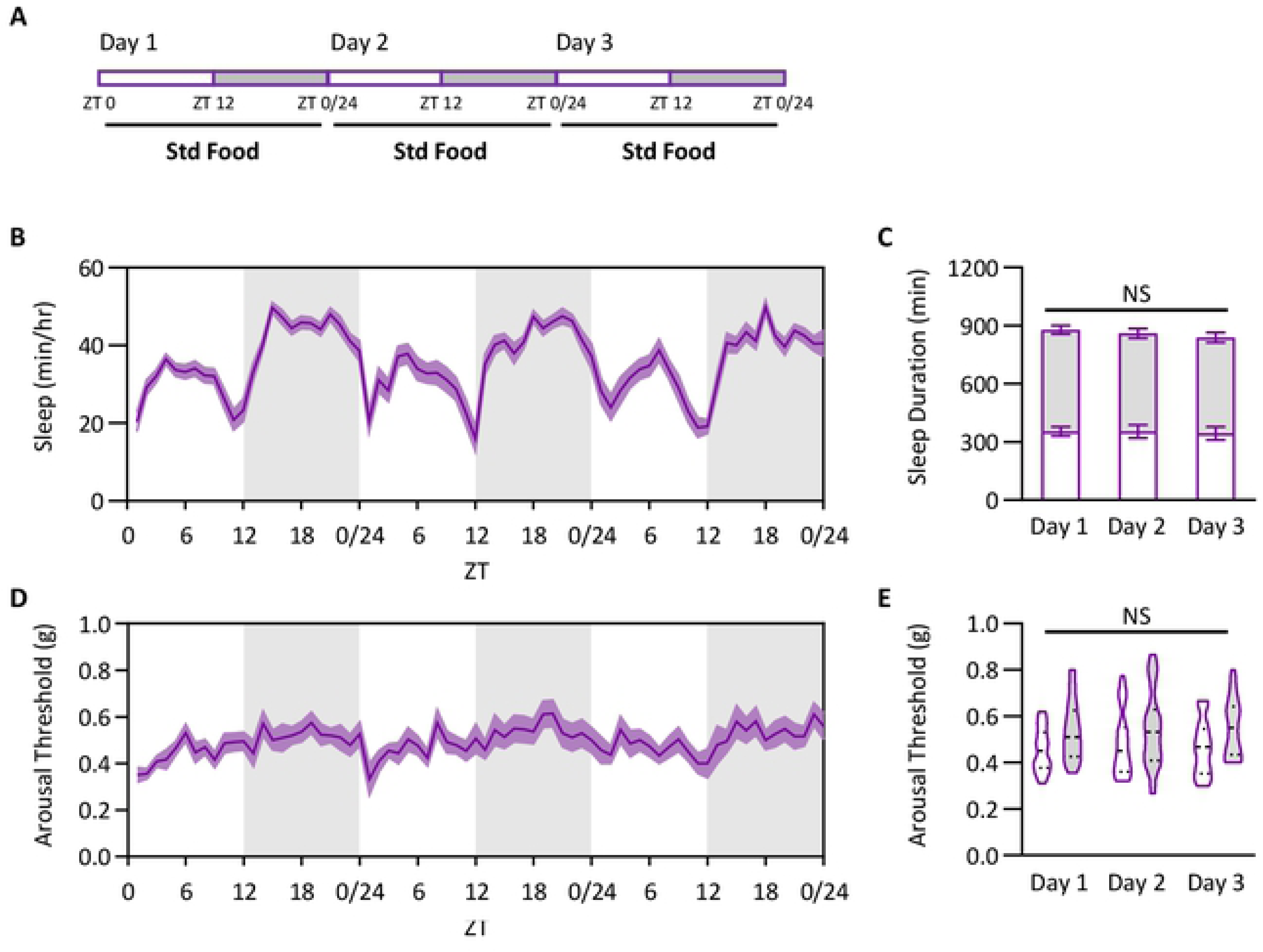
Arousal threshold does not increase as a result of acclimation to the mechanical stimulus. (A) Total sleep and arousal threshold were assessed on standard food over a 3-day period. (B) Sleep profile. (C) There is no change in sleep duration over a 3-day period (two-way ANOVA: F_2,168_ = 0.2454, *P*=0.7926). This is consistent during the daytime (ANOVA: F_2,84_ = 0.0368, *P*=0.9639) as well as the nighttime (ANOVA: F_2,84_ = 0.3558, *P*=0.7017). (D) Profile of arousal threshold. (E) There is no change in arousal threshold over a 3-day period (REML: F_2,56_ = 0.1780, *P*=0.8374). This is consistent during the daytime (Kruskal-Wallis test: H=0.0917, *P*=0.9552; N = 29) as well as the nighttime (Kruskal-Wallis test: H=0.2306, *P*=0.8911; N = 29). For sleep measurements, error bars represent +/- standard error from the mean. For arousal threshold measurements, the median (dashed line) as well as 25^th^ and 75^th^ percentiles (dotted lines) are shown. For sleep and arousal threshold profiles, shaded regions indicate +/- standard error from the mean. White background indicates daytime, while gray background indicates nighttime.

**S3 Fig.**
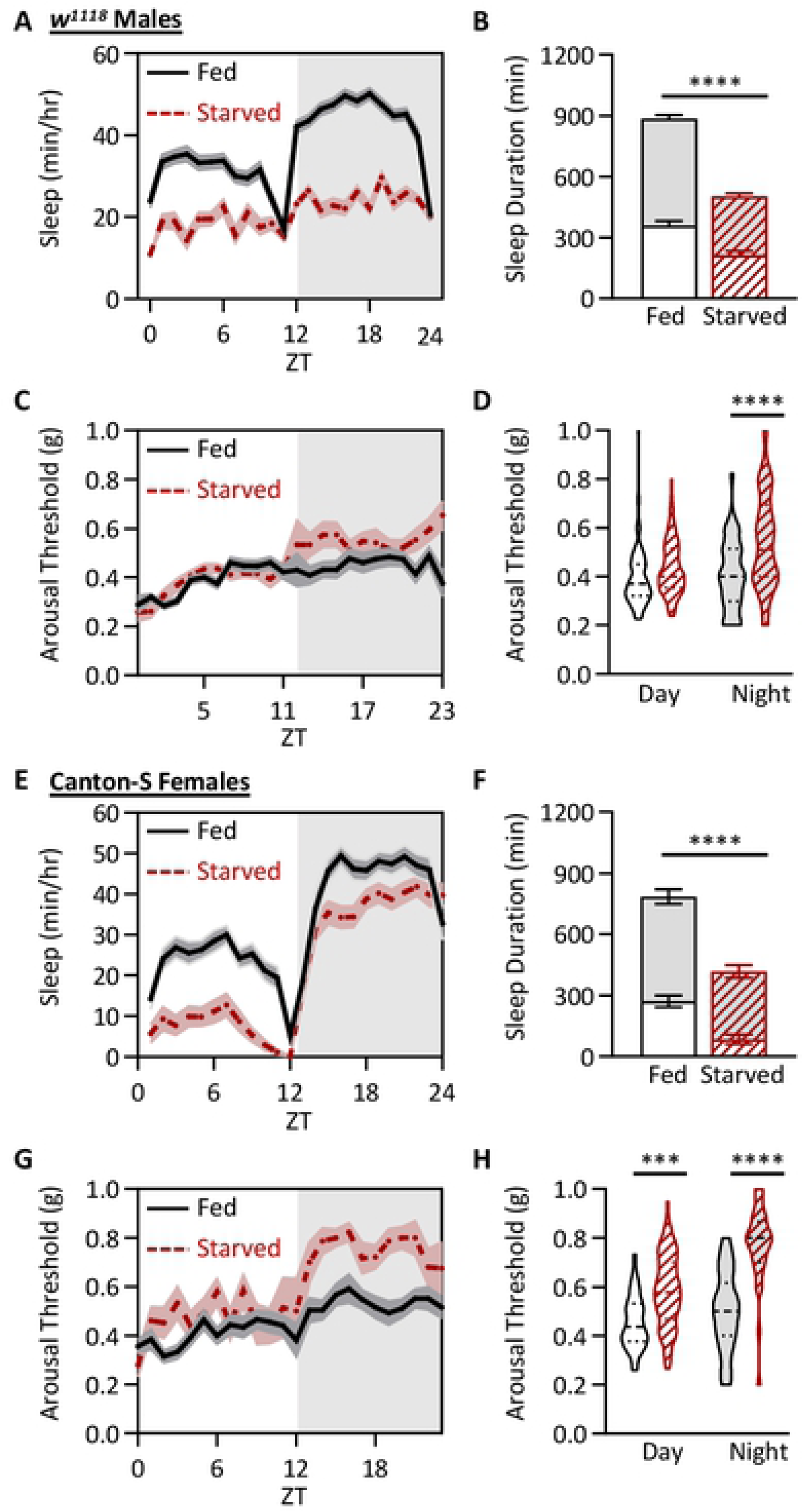
Starvation increases arousal threshold in males as well as in independent laboratory strain. All experiments were performed as described in Figure 2A. (A-D) Sleep and arousal threshold measurements in *w^1118^* male flies. (A) Sleep profiles of fed and starved *w^1118^* male flies. (B) Sleep duration decreases in the starved state (two-way ANOVA: F_1,152_ = 101.4, *P*<0.0001), and occurs in both the day (*P*<0.0001) and night (*P*<0.0001). (C) Profile of arousal threshold of fed and starved *w^1118^* male flies. (D) Arousal threshold significantly increases in the starved state (REML: F_1,71_ = 40.81, *P*<0.0001), and occurs only at night (day: *P*=0.1161; night: *P*<0.0001). (E-H) Sleep and arousal threshold measurements in female Canton-S flies. (E) Sleep profiles of fed and starved female Canton-S flies. (F) Sleep duration decreases in the starved state (two-way ANOVA: F_1,122_ = 36.92, *P*<0.0001), and occurs in both the day (*P*<0.0001) and night (*P*<0.0001). (G) Profile of arousal threshold of fed and starved female Canton-S flies. (H) Arousal threshold significantly increases in the starved state (REML: F_1,52_ = 62.11, *P*<0.0001), and occurs both during the day and at night (day: *P*=0.0008; night: *P*<0.0001). For sleep measurements, error bars represent +/- standard error from the mean. For arousal threshold measurements, the median (dashed line) as well as 25^th^ and 75^th^ percentiles (dotted lines) are shown. For sleep and arousal threshold profiles, shaded regions indicate +/- standard error from the mean. White background indicates daytime, while gray background indicates nighttime. *** = *P*<0.001; **** = *P*<0.0001.

**S4 Fig.**
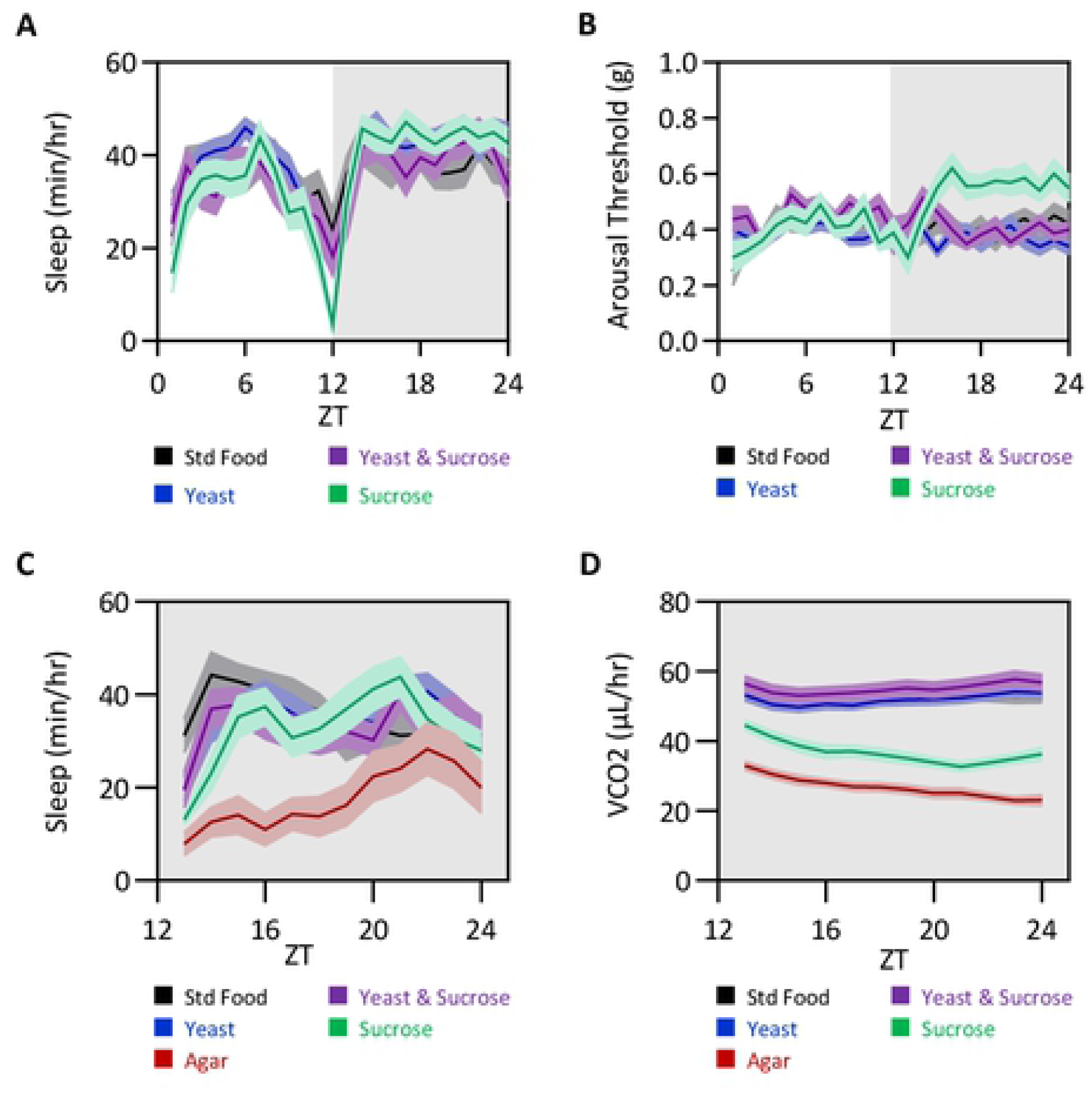
Profiles of feeding treatments in the DART and SAMM systems. Sleep, arousal threshold, and metabolic rate were taken over a 24 hr period from flies fed standard food media, 2% yeast, 2% yeast and 5% sucrose, or 5% sucrose. (A) Sleep profile. (B) Arousal threshold profile. (C) Profile of nighttime sleep duration in the SAMM system. (D) Profile of nighttime metabolic rate in the SAMM system. Shaded regions indicate +/- standard error from the mean. White background indicates daytime, while gray background indicates nighttime.

**S5 Fig.**
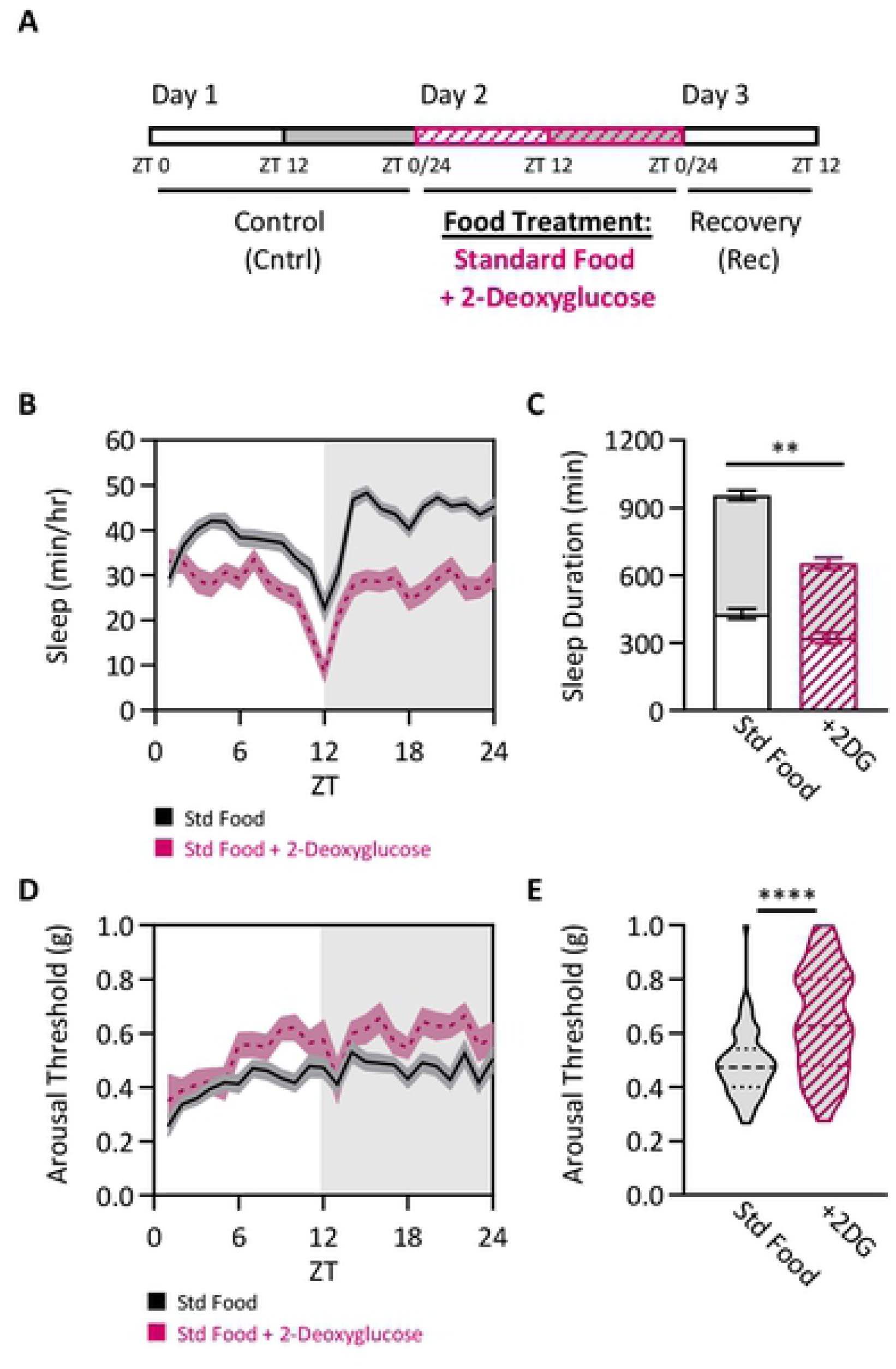
Inhibition of glycolysis phenocopies the yeast-dependent modulation of arousal threshold. (A) Total sleep and arousal threshold were assessed for 24 hrs on either standard food (black outlined boxes) or standard food + 2-deoxyglucose (+2DG; pink outlined boxes with hatches). (B) Sleep profiles of flies fed either standard food or standard + 2-deoxyglucose. (C) Sleep duration decreases in the starved state (two-way ANOVA: F_1,190_ = 38.77, *P*<0.0001), and occurs during both the day (*P*=0.0040) and night (*P*<0.0001). (D) Profile of arousal threshold of flies fed either standard food or standard + 2-deoxyglucose. (E) Nighttime arousal threshold significantly increases when 2-deoxyglucose is included in the diet (Mann-Whitney test: U=887.5, *P*<0.0001; N=46-51). For sleep measurements, error bars represent +/- standard error from mean. For arousal threshold measurements, the median (dashed line) as well as 25^th^ and 75^th^ percentiles (dotted lines) are shown. For sleep and arousal threshold profiles, shaded regions indicate +/- standard error from the mean. White background indicates daytime, while gray background indicates nighttime. ** = *P*<0.01; **** = *P*<0.0001.

**S6 Fig.**
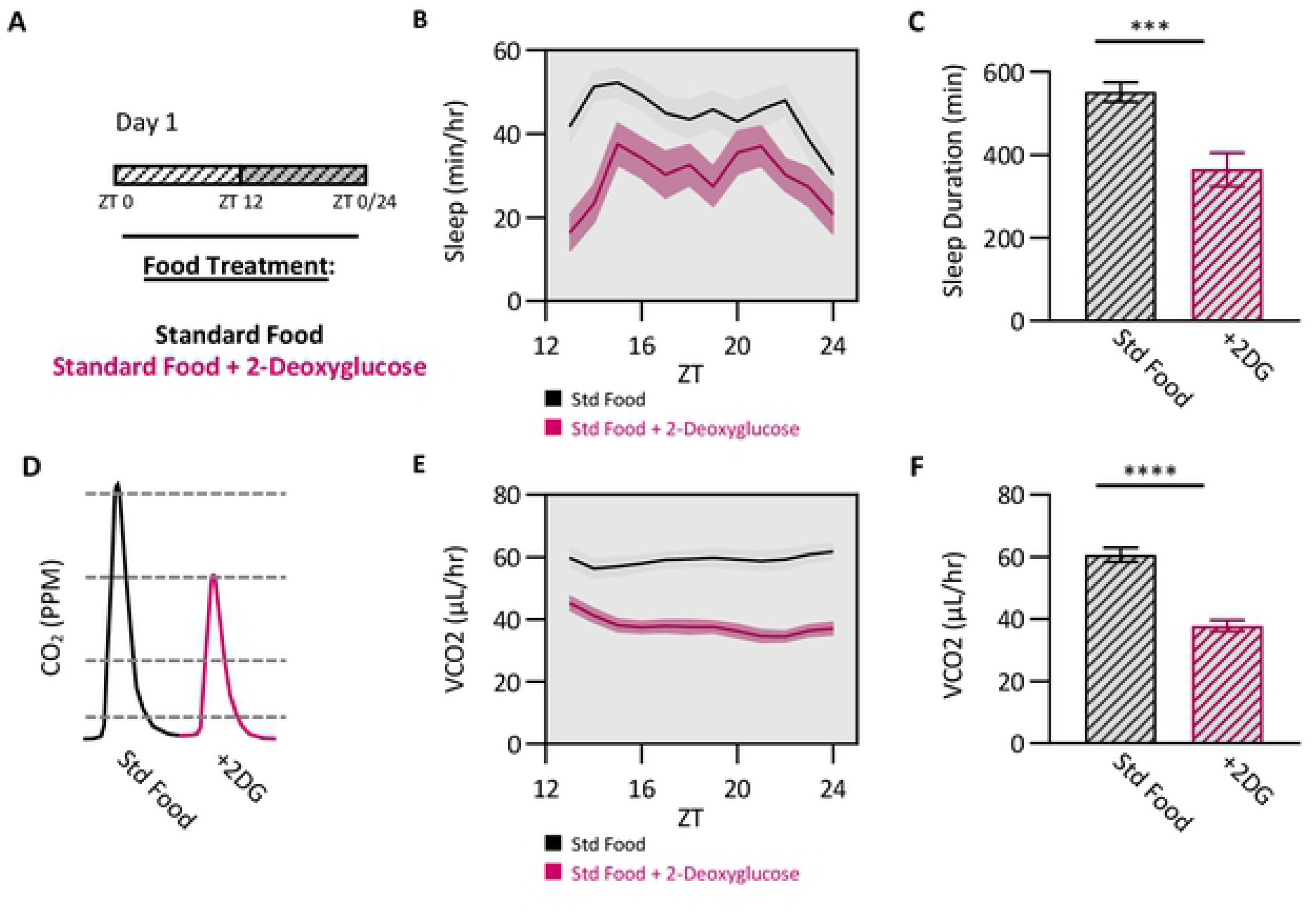
Inhibition of glycolysis phenocopies the yeast-dependent modulation of metabolic rate. (A) At ZT0, files were placed into experimental chambers containing either standard food or standard food + 2-deoxyglucose (+2DG). Files were allowed to acclimate to these conditions for 12hrs at which nighttime sleep duration and metabolic rate were then assessed. (B) Profile of nighttime sleep duration in the SAMM system. (C) Nighttime sleep duration significantly decreases in flies fed 2-deoxyglucose (t-test: t_36_=3.851, *P*=0.0005). (D) Representative traces indicating the unadjusted amount of CO_2_ produced within each experimental chamber at a given timepoint for standard food and food treated with 2-deoxyglucose. (E) Profile of nighttime metabolic rate in the SAMM system. (F) Nighttime metabolic rate significantly decreases in flies fed 2-deoxyglucose (t-test: t_36_=7.912, *P*<0.0001). Error bars represent +/- standard error from the mean. For profiles, shaded regions indicate +/- standard error from the mean. White background indicates daytime, while gray background indicates nighttime. *** = *P*<0.001; **** = *P*<0.0001.

**S7 Fig.**
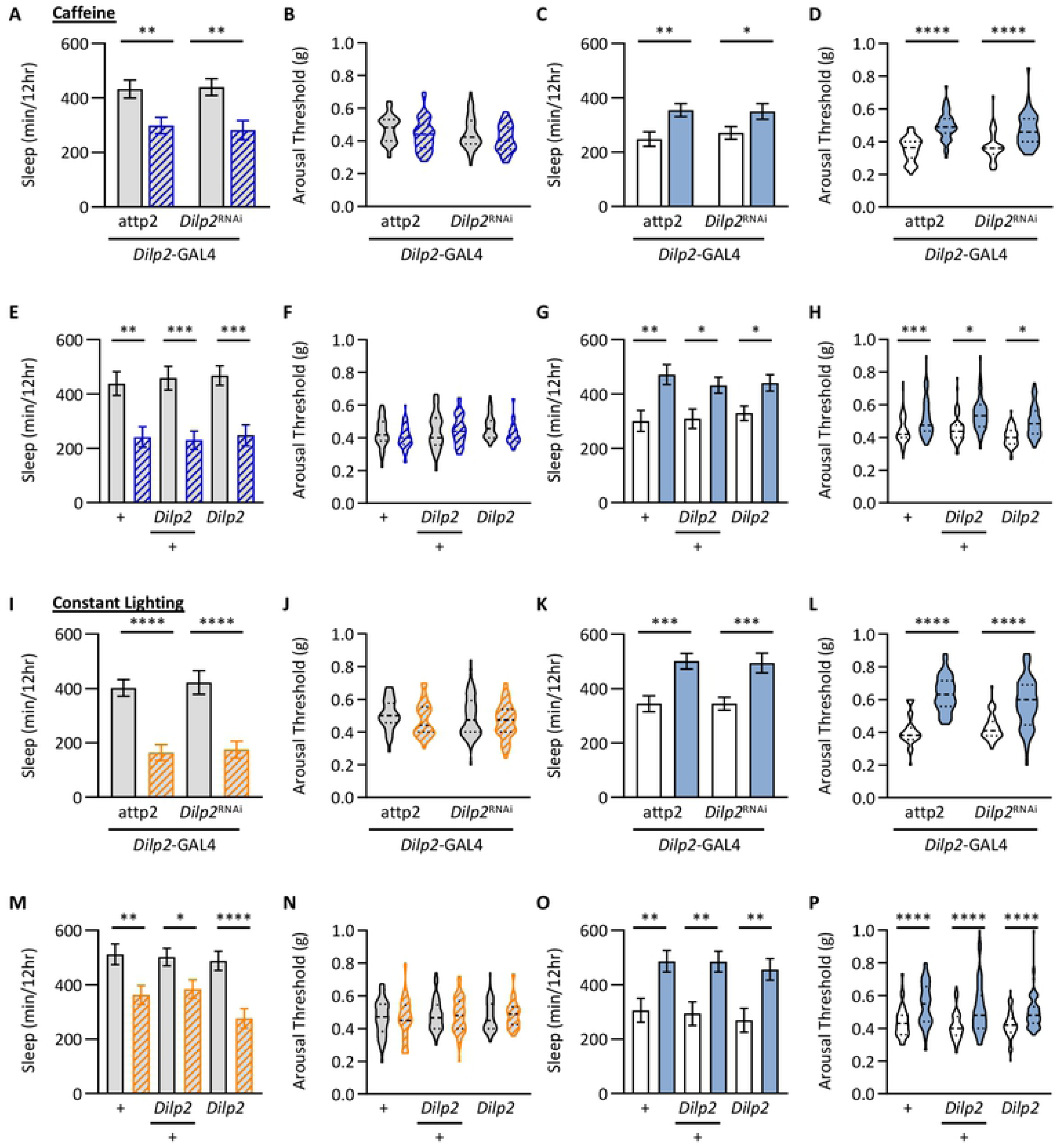
*Dilp2* does not regulate homeostatic rebound following other methods of sleep deprivation. Total sleep and arousal threshold during sleep deprivation and recovery were assessed as described in Figure 4a. However, instead of sleep depriving flies by starving them for 24 hrs, flies were sleep deprived either by adding 0.5mg/mL caffeine to their diet (A-H) or by constant lighting (I-P). (A) There is no effect of genotype on nighttime sleep duration (two-way ANOVA: F_1,154_ = 0.0240, *P*=0.8770), however there was a significant effect of being fed caffeine (two-way ANOVA: F_1,154_ = 20.56, *P*<0.0001). For each genotype, post hoc analyses revealed a significant decrease in nighttime sleep duration when fed caffeine (*Dilp*2-GAL4>UAS-*attp2*: *P*=0.0069; *Dilp*2-GAL4>UAS-*Dilp2*^RNAi^: *P*=0.0015). (B) Similar to the control (*Dilp*2-GAL4>UAS-*attp2*), knockdown of *Dilp2* in *Dilp2*-expressing neurons (*Dilp*2-GAL4>UAS-*Dilp2*^RNAi^) has no effect on nighttime arousal threshold when fed caffeine (REML: F_1,78_ = 2.227, *P*=0.1395). (C) There is no effect of genotype on daytime sleep duration (two-way ANOVA: F_1,154_ = 0.1369, *P*=0.7119), however there was a significant effect on homeostatic recovery (two-way ANOVA: F_1,154_ = 15.80, *P*=0.0001). For both genotypes, post hoc analyses revealed a significant increase in daytime sleep duration following 24 hrs of caffeine in the diet (*Dilp*2-GAL4>UAS-*attp2*: *P*=0.0028; *Dilp*2-GAL4>UAS-*Dilp2*^RNAi^: *P*=0.0371). (D) There was no effect of genotype on daytime arousal threshold upon knockdown of *Dilp2* in *Dilp2*-expressing neurons (REML: F_1,78_ = 0.0650, *P*=0.7994), however there was a significant effect on homeostatic recovery (REML: F_1,76_ = 106.1, *P*<0.0001). For all both genotypes, post hoc analyses revealed a significant increase in daytime arousal threshold following 24 hrs of caffeine in the diet (*Dilp*2-GAL4>UAS-*attp2*: *P*<0.0001; *Dilp*2-GAL4>UAS-*Dilp2*^RNAi^: *P*<0.0001). (E) There was no effect of genotype on nighttime sleep duration (two-way ANOVA: F_2,208_ = 0.1170, *P*=0.8896), however there was a significant effect of caffeine (two-way ANOVA: F_2,208_ = 45.32, *P*<0.0001). For all three genotypes, post hoc analyses revealed a significant decrease in nighttime sleep duration when caffeine was added to the diet (w^1118^: *P*=0.0016; w^1118^>*Dilp2*: *P*=0.0001; *Dilp2*: *P*=0.0003). (F) Similar to control flies (*w^1118^*), there is no change in nighttime arousal threshold when fed caffeine in *Dilp2* heterozygotes or *Dilp2* mutants (REML: F_2,105_ = 2.747, *P*=0.0686). (G) There was no effect of genotype on daytime sleep duration (two-way ANOVA: F_1,208_ = 0.1329, *P*=0.8756), however there was a significant effect on homeostatic recovery (two-way ANOVA: F_1,208_ = 25.08, *P*<0.0001). For all three genotypes, post hoc analyses revealed a significant increase in daytime sleep duration following 24 hrs of caffeine in the diet (w^1118^: *P*=0.0011; w^1118^>*Dilp2*: *P*=0.0259; *Dilp2*: *P*<0.0515). (H) There was no effect of genotype on daytime arousal threshold (REML: F_2,105_ = 0.3750, *P*<0.6880), however there was a significant effect on homeostatic recovery (REML: F_1,103_ = 29.33, *P*<0.0001). For all three genotypes, post hoc analyses revealed a significant increase in daytime arousal threshold following 24 hrs of caffeine in the diet (w^1118^: *P*=0.0004; w^1118^>*Dilp2*: *P*=0.0228; *Dilp2*: *P*=0.0218). (I) There was no effect of genotype on nighttime sleep duration (two-way ANOVA: F_1,155_ = 0.2084, *P*=0.6486), however there was a significant effect of keeping the lights on (two-way ANOVA: F_1,155_ = 50.62, *P*<0.0001). For each genotype, post hoc analyses revealed a significant decrease in nighttime sleep duration during constant lighting (*Dilp*2-GAL4>UAS-*attp2*: *P*<0.0001; *Dilp*2-GAL4>UAS-*Dilp2*^RNAi^: *P*<0.0001). (J) Similar to the control (*Dilp*2-GAL4>UAS-*attp2*), knockdown of *Dilp2* in *Dilp2*-expressing neurons (*Dilp*2-GAL4>UAS-*Dilp2*^RNAi^) has no effect on nighttime arousal threshold under constant lighting (REML: F_1,78_ = 0.4220, *P*<0.5178). (K) There was no effect of genotype on daytime sleep duration (two-way ANOVA: F_1,155_ = 0.0105, *P*=0.9184), however there was a significant effect on homeostatic recovery (two-way ANOVA: F_1,155_ = 26.64, *P*<0.0001). For all both genotypes, post hoc analyses revealed a significant increase in daytime sleep duration following 24 hrs of constant lighting (*Dilp*2-GAL4>UAS-*attp2*: *P*=0.0005; *Dilp*2-GAL4>UAS-*Dilp2*^RNAi^: *P*=0.0010). (L) There was no effect of genotype on daytime arousal threshold upon knockdown of *Dilp2* in *Dilp2*-expressing neurons (*Dilp*2-GAL4>UAS-*Dilp2*^RNAi^; REML: F_1,78_ = 0.9144, *P*=0.3418), however there was a significant effect on homeostatic recovery (REML: F_1,78_ = 157.4, *P*<0.0001). For all both genotypes, post hoc analyses revealed a significant increase in daytime arousal threshold following 24 hrs of constant lighting (*Dilp*2-GAL4>UAS-*attp2*: *P*<0.0001; *Dilp*2-GAL4>UAS-*Dilp2*^RNAi^: *P*<0.0001). (M) There was no effect of genotype on nighttime sleep duration (two-way ANOVA: F_1,229_ = 0.4038, *P*=0.6683), however there was a significant effect of keeping the lights on (two-way ANOVA: F_1,229_ = 30.98, *P*<0.0001). For all three genotypes, post hoc analyses revealed a significant decrease in nighttime sleep duration during constant lighting (w^1118^: *P*=0.0076; w^1118^>*Dilp2*: *P*=0.0026; *Dilp2*: *P*=0.0046). (N) Similar to control flies (*w^1118^*), there is no change in nighttime arousal threshold during constant lighting on in *Dilp2* heterozygotes or *Dilp2* mutants (REML: F_2,117_ = 1.652, *P*=0.1952). (O) There was no effect of genotype on sleep duration (two-way ANOVA: F_2,229_ = 0.3927, *P*=0.6757), however there was a significant effect on homeostatic recovery (two-way ANOVA: F_1,229_ = 31.02, *P*<0.0001). For all three genotypes, post hoc analyses revealed a significant increase in daytime sleep duration following 24 hrs of constant lighting (w^1118^: *P*=0.0067; w^1118^>*Dilp2*: *P*=0.0020; *Dilp2*: *P*=0.0062). (P) There was no effect of genotype on daytime arousal threshold (REML: F_2,117_ = 2.570, *P*=0.0790), however there was a significant effect on homeostatic recovery (REML: F_1,111_ = 103.8, *P*<0.0001). For all three genotypes, post hoc analyses revealed a significant increase in daytime arousal threshold following 24 hrs of caffeine in the diet (w^1118^: *P*<0.0001; w^1118^>*Dilp2*: *P*<0.0001; *Dilp2*: *P*<0.0001). For sleep measurements, error bars represent +/- standard error from the mean. For arousal threshold measurements, the median (dashed line) as well as 25^th^ and 75^th^ percentiles (dotted lines) are shown. * = *P*<0.05; ** = *P*<0.01; *** = *P*<0.001; **** = *P*<0.0001.

**S8 Fig.**
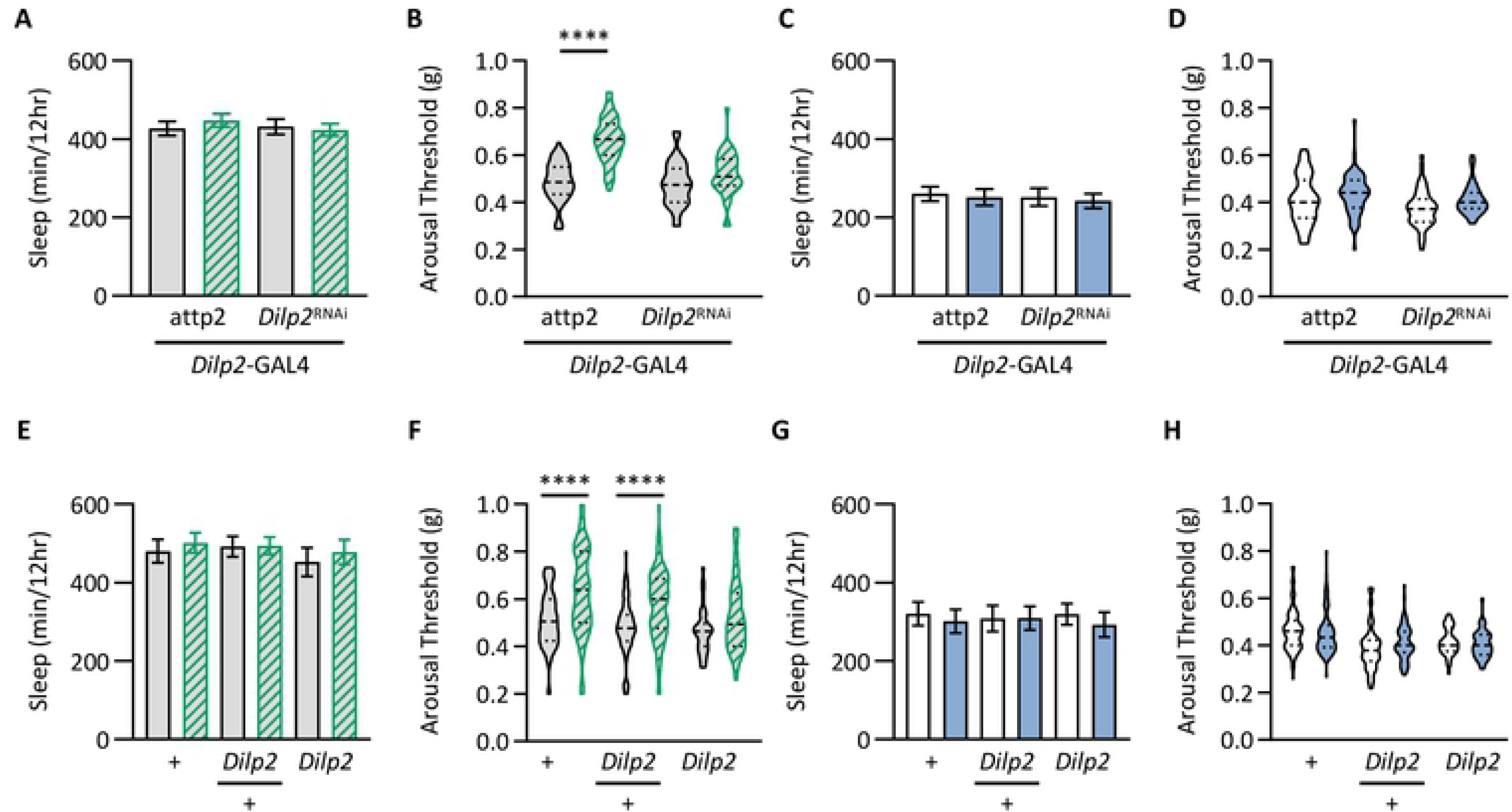
*Dilp2* uniquely regulates the yeast-dependent modulation of arousal threshold. Total sleep and arousal threshold were assessed as described in Figure 4a. However, instead of being starved for 24 hrs, flies were fed a diet of 5% sucrose. (A) Compared to the control (*Dilp*2-GAL4>UAS-*attp2*), knockdown of *Dilp2* in *Dilp2*-expressing neurons (*Dilp*2-GAL4>UAS-*Dilp2*^RNAi^) has no effect on nighttime sleep duration when fed a sucrose-only diet (two-way ANOVA: F_1,150_ = 0.3074, *P*=0.5801). (B) There is a significant effect of genotype on nighttime arousal threshold (REML: F_1,75_ = 26.51, *P*<0.0001). Post hoc analyses revealed that while controls significantly increase nighttime arousal threshold when fed a sucrose only diet (*Dilp*2-GAL4>UAS-*attp2*: *P*<0.0001), there is no effect of knockdown of *Dilp2* in *Dilp2*-expressing neurons (*Dilp*2-GAL4>UAS-*Dilp2*^RNAi^: *P*=0.1395). (C) Similar to the control (*Dilp*2-GAL4>UAS-*attp2*), knockdown of *Dilp2* in *Dilp2*-expressing neurons (*Dilp*2-GAL4>UAS-*Dilp2*^RNAi^) does not change daytime sleep duration when fed a sucrose-only diet (two-way ANOVA: F_1,150_ = 0.1921, *P*=0.6618). (D) There is a significant effect of genotype on arousal threshold following 24 hrs of starvation (REML: F_1,75_ = 6.211, *P*<0.0137). However, post hoc analyses revealed no differences in arousal threshold in the control (*Dilp*2-GAL4>UAS-*attp2*; *P*=0.2070), nor upon knockdown of *Dilp2* in *Dilp2*-expressing neurons (*Dilp*2-GAL4>UAS-*Dilp2*^RNAi^; *P*=0.1275). (E) In comparison to the control (*w^1118^*), there is no effect on nighttime sleep duration in *Dilp2* heterozygotes or *Dilp2* mutants when fed a sucrose-only diet (two-way ANOVA: F_2,216_ = 0.5651, *P*=0.5692). (F) There is a significant effect of genotype on nighttime arousal threshold (REML: F_2.108_ = 5.930, *P*=0.0032). Post hoc analyses revealed that while control flies (*w^1118^*) and *Dilp2* heterozygotes significantly increase nighttime arousal threshold when fed a diet of sucrose (w^1118^: *P*<0.0001; w^1118^>*Dilp2*: *P*<0.0001), there is no effect on arousal threshold in *Dilp2* mutants (*P*=0.1038). (G) Similar to control flies (*w^1118^*), there is no effect on daytime sleep duration following 24 hrs of a sucrose-only diet in *Dilp2* heterozygotes or *Dilp2* mutants (two-way ANOVA: F_2,216_ = 0.0130, *P*=0.9871). (H) There is a significant effect of genotype on arousal threshold following 24 hrs of starvation (REML: F_2,108_ = 14.58, *P*<0.0001). However, post hoc analyses revealed no differences in arousal threshold in control flies (*w^1118^*: *P*=0.6729), *Dilp2* heterozygotes (*w^1118^*>*Dilp2*: *P*=0.14111), nor *Dilp2* mutants (*Dilp*2: *P*=0.9464). For sleep measurements, error bars represent +/- standard error from the mean. For arousal threshold measurements, the median (dashed line) as well as 25^th^ and 75^th^ percentiles (dotted lines) are shown. **** = *P*<0.0001.

**S9 Fig.**
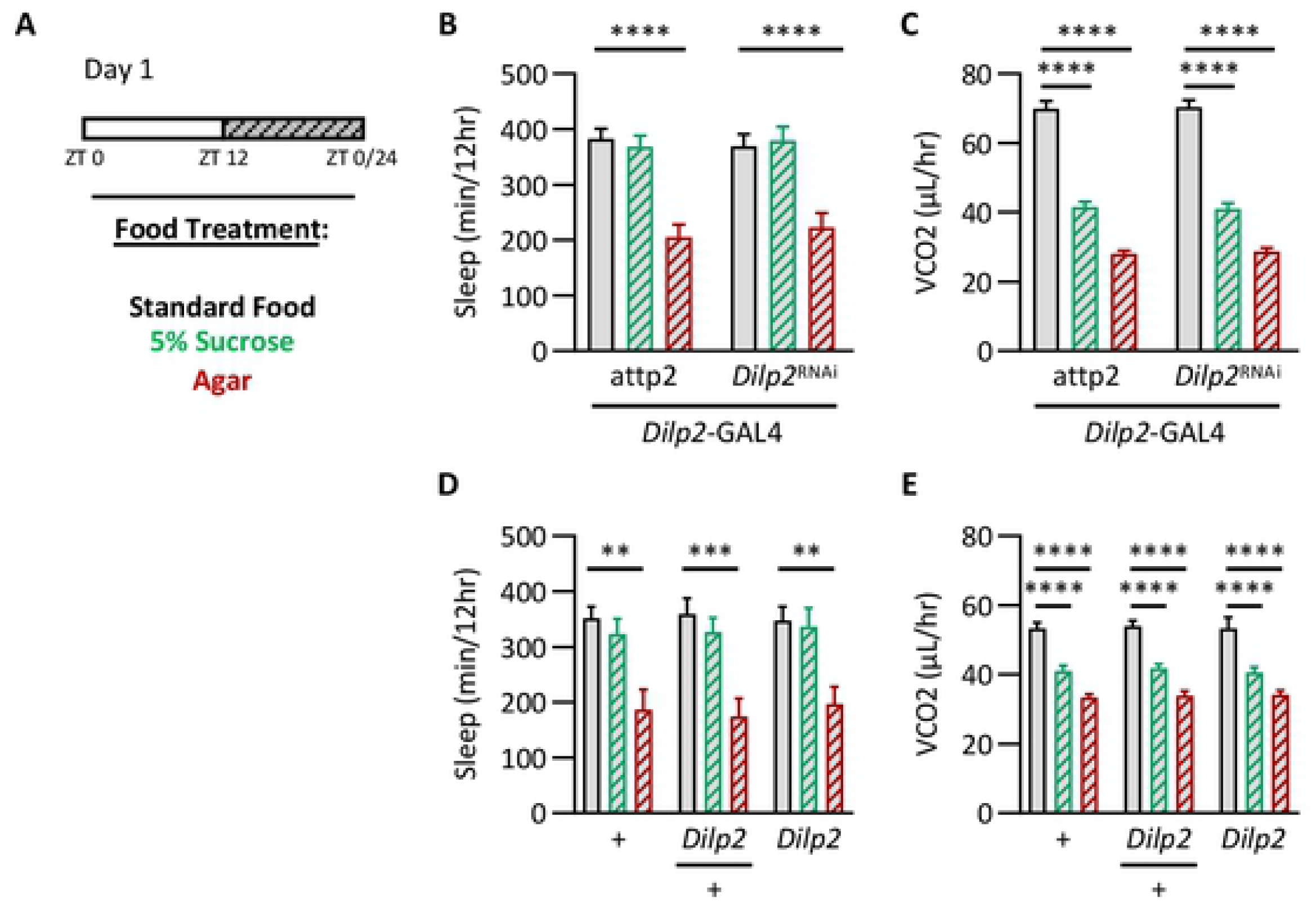
*Dilp2* does not regulate metabolic rate. Nighttime sleep duration and metabolic rate were assessed in the SAMM system as described in Figure 3e. (A) Flies were fed a diet of either standard food, sucrose, or agar. (B) Compared to the control (*Dilp*2-GAL4>UAS-*attp2*), knockdown of *Dilp2* in *Dilp2*-expressing neurons (*Dilp*2-GAL4>UAS-*Dilp2*^RNAi^) has no effect on nighttime sleep duration (two-way ANOVA: F_1,137_ = 0.07386, *P*=0.7862), however there was a significant effect of diet (two-way ANOVA: F_2,137_ = 33.67, *P*<0.0001). For each genotype, post hoc analyses revealed a significant decrease in nighttime sleep duration when starved (*Dilp*2-GAL4>UAS-*attp2*: *P*<0.0001; *Dilp*2-GAL4>UAS-*Dilp2*^RNAi^: *P*<0.0001), when compared to standard food. (C) Compared to the control (*Dilp*2-GAL4>UAS-*attp2*), knockdown of *Dilp2* in *Dilp2*-expressing neurons (*Dilp*2-GAL4>UAS-*Dilp2*^RNAi^) has no effect on metabolic rate (two-way ANOVA: F_1,137_ = 330.9, *P*=0.8969), however there was a significant effect of diet (two-way ANOVA: F_2,137_ = 330.9, *P*<0.0001). For each genotype, post hoc analyses revealed a significant decrease in metabolic when fed a sucrose-only diet (*Dilp*2-GAL4>UAS-*attp2*: *P*<0.0001; *Dilp*2-GAL4>UAS-*Dilp2*^RNAi^: *P*<0.0001) as well as when starved (*Dilp*2-GAL4>UAS-*attp2*: *P*<0.0001; *Dilp*2-GAL4>UAS-*Dilp2*^RNAi^: *P*<0.0001), when compared to standard food. (D) In comparison to the control (*w^1118^*), there is no effect on nighttime sleep duration in *Dilp2*^null^ heterozygotes or *Dilp2*^null^ mutants (two-way ANOVA: F_2,145_ = 0.1374, *P*<0.8717), however there was a significant effect of diet (two-way ANOVA: F_2,145_ = 24.77, *P*<0.0001). For all three genotypes, post hoc analyses revealed a significant decrease in nighttime sleep duration when starved (w^1118^: *P*=0.0011; w^1118^>*Dilp2*^null^: *P*=0.0002; *Dilp2*^null^: *P*=0.0038). (E) In comparison to the control (*w^1118^*), there is no effect on metabolic rate in *Dilp2*^null^ heterozygotes or *Dilp2*^null^ mutants (two-way ANOVA: F_2,164_ = 0.1528, *P*=0.8584), however there was a significant effect of diet (two-way ANOVA: F_2,164_ = 120.3, *P*<0.0001). For all three genotypes, post hoc analyses revealed a significant decrease in metabolic when fed a sucrose-only diet (w^1118^: *P*<0.0001; w^1118^>*Dilp2*^null^: *P*<0.0001; *Dilp2*^null^: *P*<0.0001) as well as when starved (w^1118^: *P*<0.0001; w^1118^>*Dilp2*^null^: *P*<0.0001; *Dilp2*^null^: *P*<0.0001), when compared to standard food. Error bars represent +/- standard error from the mean. ** = *P*<0.01; *** = *P*<0.001; **** = *P*<0.0001.

## References

1. Keene AC, Duboue ER. The origins and evolution of sleep. J Exp Biol. 2018; doi:10.1242/jeb.159533

2. Joiner WJ. Unraveling the Evolutionary Determinants of Sleep. Curr Biol. Elsevier Ltd; 2016;26: R1073–R1087. doi:10.1016/j.cub.2016.08.068

3. Allada R, Siegel JM. Unearthing the phylogenetic roots of sleep. Curr Biol. 2008;18: R670–R679. doi:10.1016/j.cub.2008.06.033

4. Donlea JM. Neuronal and molecular mechanisms of sleep homeostasis. Curr Opin Insect Sci. Elsevier Inc; 2017;24: 51–57. doi:10.1016/j.cois.2017.09.008

5. Tougeron K, Abram PK. An Ecological Perspective on Sleep Disruption. Am Nat. 2017;190: E55–E66. doi:10.1086/692604

6. Campbell SS, Tobler I. Animal sleep: a review of sleep duration across phylogeny. Neurosci Biobehav Rev. 1984;8: 269–300. doi:10.1016/0149-7634(84)90054-X

7. Hartmann E. The function of sleep. Annu Psychoanal. 1974;2: 271–289.

8. Seidner G, Robinson J, Wu M, Worden K, Masek P, Roberts S, et al. Identification of Neurons with a Privileged Role in Sleep Homeostasis in *Drosophila melanogaster*. Curr Biol. 2015;25: 2928–39.

9. Liu S, Liu Q, Tabuchi M, Wu M. Sleep Drive Is Encoded by Neural Plastic Changes in a Dedicated Circuit. Cell. 2016;165: 1347–60.

10. Danguir J, Nicolaidis S. Dependence of sleep on nutrient’s availability. Physiology and Behavior. 1979. pp. 735–740. doi:10.1016/0031-9384(79)90240-3

11. Keene AC, Duboué ER, McDonald DM, Dus M, Suh GSB, Waddell S, et al. Clock and cycle limit starvation-induced sleep loss in *Drosophila*. Curr Biol. 2010;20: 1209–1215. doi:10.1016/j.cub.2010.05.029

12. Griffith LC. Neuromodulatory control of sleep in *Drosophila melanogaster*: Integration of competing and complementary behaviors. Curr Opin Neurobiol. Elsevier Ltd; 2013;23: 819–823. doi:10.1016/j.conb.2013.05.003

13. Yurgel M, Masek P, DiAngelo JR, Keene A. Genetic dissection of sleep-metabolism interactions in the fruit fly. J Comp Physiol A Neuroethol Sens Neural Behav Physiol. 2014;epub ahead.

14. Crocker A, Shahidullah M, Levitan IB, Sehgal A. Identification of a Neural Circuit that Underlies the Effects of Octopamine on Sleep:Wake Behavior. Neuron. Elsevier Inc.; 2010;65: 670–681. doi:10.1016/j.neuron.2010.01.032

15. Kreneisz O, Chen X, Fridell YW, Mulkey DK. Glucose increases activity and Ca2+ in insulin-producing cells of adult *Drosophila*. Neuroreport. 2010;21: 1116–1120. doi:10.1097/WNR.0b013e3283409200

16. Varin C, Rancillac A, Geoffroy H, Arthaud S, Fort P, Gallopin T. Glucose Induces Slow-Wave Sleep by Exciting the Sleep-Promoting Neurons in the Ventrolateral Preoptic Nucleus: A New Link between Sleep and Metabolism. J Neurosci. 2015;35: 9900–9911. doi:10.1523/JNEUROSCI.0609-15.2015

17. Chakravarti L, Moscato EH, Kayser MS. Unraveling the Neurobiology of Sleep and Sleep Disorders Using Drosophila. 2017;121: 253–285. doi:10.1016/bs.ctdb.2016.07.010

18. Artiushin G, Sehgal A. The *Drosophila* circuitry of sleep–wake regulation. Curr Opin Neurobiol. 2017;44: 243–250. doi:10.1016/j.conb.2017.03.004

19. Donlea JM, Pimentel D, Talbot CB, Kempf A, Omoto JJ, Hartenstein V, et al. Recurrent Circuitry for Balancing Sleep Need and Sleep. Neuron. 2018; 1–12. doi:10.1016/j.neuron.2017.12.016

20. Pimentel D, Donlea JM, Talbot CB, Song SM, Thurston AJF, Miesenböck G. Operation of a homeostatic sleep switch. Nature. Nature Publishing Group; 2016;536: 333–337. doi:10.1038/nature19055

21. Toda H, Williams JA, Gulledge M, Sehgal A. A sleep-inducing gene, nemuri, links sleep and immune function in *Drosophila*. Science (80-). 2019;363: 509–515. doi:10.1126/science.aat1650

22. Thimgan MS, Suzuki Y, Seugnet L, Gottschalk L, Shaw PJ. The Perilipin Homologue, Lipid Storage Droplet 2, Regulates Sleep Homeostasis and Prevents Learning Impairments Following Sleep Loss. 2010;8. doi:10.1371/journal.pbio.1000466

23. Donlea J, Leahy A, Thimgan MS, Suzuki Y, Hughson BN, Sokolowski MB, et al. foraging alters resilience/vulnerability to sleep disruption and starvation in *Drosophila*. Proc Natl Acad Sci. 2012;109: 2613–2618. doi:10.1073/pnas.1112623109

24. Alphen B van, Yap MHW, Kirszenblat L, Kottler B, Swinderen B van. A Dynamic Deep Sleep Stage in *Drosophila*. J Neurosci. 2013;33: 6917–6927. doi:10.1523/JNEUROSCI.0061-13.2013

25. Faville R, Kottler B, Goodhill G, Shaw PJ, van Swinderen B. How deeply does your mutant sleep? Probing arousal to better understand sleep defects in *Drosophila*. Sci Reports. 2015;13: 8454.

26. Yap MHW, Grabowska MJ, Rohrscheib C, Jeans R, Troup M, Paulk AC, et al. Oscillatory brain activity in spontaneous and induced sleep stages in flies. Nat Commun. Springer US; 2017;8. doi:10.1038/s41467-017-02024-y

27. Stahl BA, Slocumb ME, Chaitin H, DiAngelo JR, Keene AC. Sleep-Dependent Modulation of Metabolic Rate in *Drosophila*. Sleep. 2017;40. doi:10.1093/sleep/zsx084

28. Harshman LG, Hoffmann AA, Clark AG. Selection for starvation resistance in *Drosophila melanogaster*: Physiological correlates, enzyme activities and multiple stress responses. J Evol Biol. 1999;12: 370–379. doi:10.1046/j.1420-9101.1999.00024.x

29. Andretic R, Shaw PJ. Essentials of sleep recordings in *Drosophila*: Moving beyond sleep time. Methods in Enzymology. 2005. pp. 759–772. doi:10.1016/S0076-6879(05)93040-1

30. Brown EB, Torres J, Bennick RA, Rozzo V, Kerbs A, Diangelo JR, et al. Variation in sleep and metabolic function is associated with latitude and average temperature in *Drosophila melanogaster*. Ecol Evol. 2018; doi:10.1002/ece3.3963

31. Khericha M, Kolenchery JB, Tauber E. Neural and non-neural contributions to sexual dimorphism of mid-day sleep in *Drosophila melanogaster*: a pilot study. Physiol Entomol. 2016;41: 327–334. doi:10.1111/phen.12134

32. Kempf A, Song SM, Talbot CB, Miesenböck G. A potassium channel β-subunit couples mitochondrial electron transport to sleep. Nature. 2019;568: 230–234. doi:10.1038/s41586-019-1034-5

33. De Camargo R, Phaff HJ. YEASTS OCCURRING IN *DROSOPHILA* FLIES AND IN FERMENTING TOMATO FRUITS IN NORTHERN CALIFORNIA. J Food Sci. 1957;22: 367–372. doi:10.1111/j.1365-2621.1957.tb17024.x

34. Phaff HJ, Miller MW, Recca JA, Shifrine M, Mrak EM. Yeasts Found in the Alimentary Canal of *Drosophila*. Ecology. John Wiley & Sons, Ltd; 1956;37: 533–538. doi:10.2307/1930176

35. Libert S, Zwiener J, Chu X, VanVoorhies W, Roman G, Pletcher SD. Regulation of *Drosophila* life span by olfaction and food-derived odors. Science (80-). 2007;315: 1133– 7. doi:10.1126/science.1136610

36. Linford NJ, Chan TP, Pletcher SD. Re-patterning sleep architecture in *Drosophila* through gustatory perception and nutritional quality. PLoS Genet. 2012;8. doi:10.1371/journal.pgen.1002668

37. Dus M, Min S, Keene AC, Lee GY, Suh GSB. Taste-independent detection of the caloric content of sugar in *Drosophila*. Proc Natl Acad Sci U S A. 2011;108: 11644–9. doi:10.1073/pnas.1017096108

38. Murakami K, Yurgel ME, Stahl BA, Masek P, Mehta A, Heidker R, et al. Translin Is Required for Metabolic Regulation of Sleep. Curr Biol. Elsevier Ltd; 2016;26: 972–980. doi:10.1016/j.cub.2016.02.013

39. Stahl B, Slocumb M, Chaitin H, DiAngelo J, Keene A. Sleep-Dependent Modulation Of Metabolic Rate In *Drosophila*. Sleep. 2017;in press.

40. DiAngelo JR, Erion R, Crocker A, Sehgal A. The central clock neurons regulate lipid storage in *Drosophila*. PLoS One. 2011;6. doi:10.1371/journal.pone.0019921

41. Rajan A, Perrimon N. *Drosophila* Cytokine Unpaired 2 Regulates Physiological Homeostasis by Remotely Controlling Insulin Secretion. Cell. 2012. pp. 123–137. doi:10.1016/j.cell.2012.08.019

42. Ikeya T, Galic M, Belawat P, Nairz K, Hafen E. Nutrient-dependent expression of insulin-like peptides from neuroendocrine cells in the CNS contributes to growth regulation in *Drosophila*. Curr Biol. 2002;12: 1293–1300. doi:10.1016/S0960-9822(02)01043-6

43. Birse RT, Choi J, Reardon K, Rodriguez J, Graham S, Diop S, et al. High-fat-diet-induced obesity and heart dysfunction are regulated by the TOR pathway in *Drosophila*. Cell Metab. 2010;12: 533–544. doi:10.1016/j.cmet.2010.09.014

44. Metaxakis A, Tain L, Grönke, Hendrich O, Birras U, Partridge L. Lowered insulin signalling ameliorates age-related sleep fragmentation in *Drosophila*. PLoS Biol. 2014;12: e1001824.

45. Cong X, Wang H, Liu Z, An C, Zhao Z. Regulation of Sleep by Insulin-like Peptide System in *Drosophila melanogaster*. Sleep. 2015; sp-00236-14.

46. Grönke S, Clarke D-F, Broughton S, Andrews TD, Partridge L. Molecular evolution and functional characterization of *Drosophila* insulin-like peptides. PLoS Genet. 2010;6: e1000857. doi:10.1371/journal.pgen.1000857

47. Nath RD, Bedbrook CN, Abrams MJ, Basinger T, Bois JS, Prober DA, et al. The Jellyfish Cassiopea Exhibits a Sleep-like State. Curr Biol. 2017;27: 2984–2990.e3. doi:10.1016/j.cub.2017.08.014

48. Krashes MJ, DasGupta S, Vreede A, White B, Armstrong JD, Waddell S. A Neural Circuit Mechanism Integrating Motivational State with Memory Expression in *Drosophila*. Cell. 2009;139: 416–27. doi:10.1016/j.cell.2009.08.035

49. Krashes MJ, Waddell S. Rapid consolidation to a radish and protein synthesis-dependent long-term memory after single-session appetitive olfactory conditioning in *Drosophila*. J Neurosci. 2008;19: 3103–13. doi:10.1523/JNEUROSCI.5333-07.2008

50. Cervantes-Sandoval I, Davis RL. Distinct traces for appetitive versus aversive olfactory memories in DPM neurons of *Drosophila*. Curr Biol. 2012;22: 1247–52. doi:10.1016/j.cub.2012.05.009

51. Borbély AA. A two process model of sleep regulation. Hum Neurobiol. 1982;1: 195–204. doi:10.1111/jsr.12371

52. Donlea JM, Pimentel D, Miesenböck G. Neuronal machinery of sleep homeostasis in *Drosophila*. Neuron. 2014;81: 860–872. doi:10.1016/j.neuron.2013.12.013

53. Griffith LC, Hobin M, Van Vactor D, Fulga TA, Moore J, Goodwin PR, et al. MicroRNAs Regulate Sleep and Sleep Homeostasis in *Drosophila*. Cell Rep. ElsevierCompany.; 2018;23: 3776–3786. doi:10.1016/j.celrep.2018.05.078

54. Gerstner JR, Taylor RH, Frank MG, Van Dongen HPA, Vanderheyden WM, Goodman AG. Astrocyte expression of the *Drosophila* TNF-alpha homologue, Eiger, regulates sleep in flies. PLOS Genet. 2018;14: e1007724. doi:10.1371/journal.pgen.1007724

55. Allada R, Cirelli C, Sehgal A. Molecular Mechanisms of Sleep Homeostasis in Flies and Mammals. Cold Spring Harb Perspect Biol. 2017; a027730. doi:10.1101/cshperspect.a027730

56. Liu G, Seiler H, Wen A, Zars T, Ito K, Wolf R, et al. Distinct memory traces for two visual features in the *Drosophila* brain. Nature. 2006;439: 551–6. doi:10.1038/nature04381

57. Pan Y, Zhou Y, Guo C, Gong H, Gong Z, Liu L. Differential roles of the fan-shaped body and the ellipsoid body in *Drosophila* visual pattern memory. 2009; 289–295. doi:10.1101/lm.1331809.16

58. Sitaraman D, Aso Y, Jin X, Chen N, Felix M, Rubin GM, et al. Propagation of Homeostatic Sleep Signals by Segregated Synaptic Microcircuits of the *Drosophila* Mushroom Body. Curr Biol. 2015;25: 2517–2527. doi:10.1016/j.cub.2015.09.017

59. Tabuchi M, Lone SR, Liu Q, Zhang J, Spira A, Wu M. Sleep Interacts with Abeta to Modulate Intrinsic Neuronal Excitability. Curr Bilogy.

60. Shaw PJ, Cirelli C, Greenspan RJ, Tononi G. Correlates of sleep and waking in *Drosophila melanogaster*. Science. 2000;287: 1834–1837. doi:10.1126/science.287.5459.1834

61. Beckwith E, Geissmann Q, French A, Gilestro G. Regulation of sleep homeostasis by sexual arousal. Elife. 2017;6: e27445.

62. Machado DR, Afonso DJS, Kenny AR, Öztürk-Çolak A, Moscato EH, Mainwaring B, et al. Identification of octopaminergic neurons that modulate sleep suppression by male sex drive. Elife. 2017;6: 1–21. doi:10.7554/eLife.23130

63. Szymusiak R. Hypothalamic versus neocortical control of sleep. Curr Opin Pulm Med. 2010;16: 530–5. doi:10.1097/MCP.0b013e32833eec92

64. Hanlon EC, Vyazovskiy V V, Faraguna U, Tononi G, Cirelli C. Synaptic Potentiation and Sleep Need : Clues from Molecular and Electrophysiological Studies. Curr Top Med Chem. 2011;11: 2472–2482. doi:BSP/CTMC/E-Pub/-000191-11-21 [pii]

65. Hasegawa T, Tomita J, Hashimoto R, Ueno T, Kume S, Kume K. Sweetness induces sleep through gustatory signalling independent of nutritional value in a starved fruit fly. Sci Rep. Springer US; 2017;7: 1–9. doi:10.1038/s41598-017-14608-1

66. Sonn JY, Lee J, Sung MK, Ri H, Choi JK, Lim C, et al. Serine metabolism in the brain regulates starvation-induced sleep suppression in *Drosophila melanogaster*. Proc Natl Acad Sci U S A. 2018; 201719033. doi:10.1073/pnas.1719033115

67. Yurgel ME, Kakad P, Zandawala M, Nässel DR, Godenschwege TA, Keene AC. A single pair of leucokinin neurons are modulated by feeding state and regulate sleep–metabolism interactions. PLoS Biol. 2019;17: e2006409. doi:10.1371/journal.pbio.2006409

68. Keene AC, Duboué ER, McDonald DM, Dus M, Suh GSB, Waddell S, et al. Clock and cycle limit starvation-induced sleep loss in *Drosophila*. Curr Biol. 2010;20. doi:10.1016/j.cub.2010.05.029

69. Linford NJ, Ro J, Chung BY, Pletcher SD. Gustatory and metabolic perception of nutrient stress in *Drosophila*. Proc Natl Acad Sci U S A. 2015;112: 2587–92. doi:10.1073/pnas.1401501112

70. Broughton SJ, Slack C, Alic N, Metaxakis A, Bass TM, Driege Y, et al. DILP-producing median neurosecretory cells in the *Drosophila* brain mediate the response of lifespan to nutrition. Aging Cell. 2010;9: 336–346. doi:10.1111/j.1474-9726.2010.00558.x

71. Cong X, Wang H, Liu Z, He C, An C, Zhao Z. Regulation of Sleep by Insulin-like Peptide System in *Drosophila melanogaster*. Sleep. 2015;38: 1075–83. doi:10.5665/sleep.4816

72. Rulifson EJ, Kim SK, Nusse R. Insulin-Producing Neurons in Flies : Growth and Diabetic Phenotypes. Science (80-). 2002;296: 1118–1120.

73. Ikeya T, Broughton S, Vinti G, Bass T, Alic N, Partridge L, et al. Reduction of DILP2 in *Drosophila* Triages a Metabolic Phenotype from Lifespan Revealing Redundancy and Compensation among DILPs. PLoS One. 2008;3: e3721. doi:10.1371/journal.pone.0003721

74. Post S, Karashchuk G, Wade JD, Sajid W, De Meyts P, Tatar M. *Drosophila* insulin-like peptides DILP2 and DILP5 differentially stimulate cell signaling and glycogen phosphorylase to regulate longevity. Front Endocrinol (Lausanne). 2018;9: 1–16. doi:10.3389/fendo.2018.00245

75. Birse RT, Soderberg JAE, Luo J, Winther AME, Nassel DR. Regulation of insulin-producing cells in the adult *Drosophila* brain via the tachykinin peptide receptor DTKR. J Exp Biol. 2011;214: 4201–4208. doi:10.1242/jeb.062091

76. Sudhakar SR, Varghese J. Insulin signalling activates multiple feedback loops to elicit hunger-induced feeding in *Drosophila*. Abstract: 2018; doi:10.1101/364554

77. Chung BY, Ro J, Hutter SA, Miller KM, Guduguntla LS, Kondo S, et al. *Drosophila* Neuropeptide F Signaling Independently Regulates Feeding and Sleep-Wake Behavior. Cell Rep. ElsevierCompany.; 2017;19: 2441–2450. doi:10.1016/j.celrep.2017.05.085

78. Ni J, Markstein M, Binari R, Pfeiffer B, Liu L, Villalta C, et al. Vector and parameters for targeted transgenic RNA interference in *Drosophila melanogaster*. Nat Methods. 2008;5: 49–51. doi:10.1038/nmeth1146

79. Kubrak OI, Lushchak O V., Zandawala M, Nässel DR. Systemic corazonin signalling modulates stress responses and metabolism in *Drosophila*. Open Biol. 2016;6: 160152. doi:10.1098/rsob.160152

